# Uncharacterized unicellular relatives reveal key gaps in sampling diversity for reconstructing animal origins

**DOI:** 10.64898/2026.10.06.756977

**Authors:** Alicia Jiménez-Valadez, Iñaki Ruiz-Trillo, Jennah E. Dharamshi, Cédric Berney

## Abstract

The origin of multicellular animals from unicellular ancestors remains a major evolutionary question. To address this transition, a reconstruction of the nature of the last unicellular ancestor of animals (LUAA) is key, and requires broad sampling of extant unicellular relatives, which together with animals form the clade Holozoa. Environmental surveys suggest that unicellular Holozoa diversity extends beyond the five groups with characterized members (Choanoflagellatea, Ichthyosporea, Filasterea, Corallochytrea, and Atreyea). However, the extent of the environmental diversity within and outside these groups remains unclear. To fill this gap, we here systematically re-evaluated Holozoa diversity using environmental 18S rDNA sequences compiled from updated datasets, including long-read metabarcoding surveys. We report nine uncharacterized lineages inside Holozoa (seven newly identified here), provisionally placed outside the described groups. We also uncovered a previously unreported diverse lineage deeper inside Opisthokonta, separate from Holozoa and Fungi. In addition, we identified 21 uncharacterized lineages within Choanoflagellatea and Ichthyosporea (eleven newly reported here). We used short-read metabarcoding datasets to characterize the environmental and geographical distributions of these lineages. Many are globally distributed and some can reach relative abundances comparable to those of the described groups, while others are restricted to specific environments. Our findings reveal substantial uncharacterized diversity missing from the cultured taxa used in comparative studies of animal origins. By placing this diversity in a phylogenetic and environmental context, our study will guide the targeted isolation efforts needed to broaden the taxon sampling available to reconstruct LUAA and unravel the unicellular-to-multicellular transition that gave rise to animals.

## Introduction

Unicellular-to-multicellular transitions are among the major evolutionary innovations in the history of life. Among these, the origin of animals (Metazoa) has long attracted particular attention due to the morphological diversity and developmental complexity of animals, and our own membership in the group (King 2004; Ruiz-Trillo et al. 2007; Butterfield 2011; Erwin et al. 2011; Brunet and King 2017; Ruiz-Trillo et al. 2023). Yet, the nature of the unicellular ancestor from which animals arose remains unknown. Reconstructing the origin of animals requires understanding both their last common ancestor (LACA) and its unicellular predecessor, the last unicellular animal ancestor (LUAA) (Ros-Rocher et al. 2021; Ruiz-Trillo et al. 2026). Whereas LACA can be inferred through comparative analyses of extant animals, reconstructing LUAA requires broad sampling across the diversity of unicellular relatives (Suga et al. 2012; Richter et al. 2018; Ocaña-Pallarès et al. 2022; Ruiz-Trillo et al. 2023).

Metazoa and their unicellular relatives comprise the clade Holozoa (Lang et al. 2002; Ruiz-Trillo et al. 2008; Shalchian-Tabrizi et al. 2008; Torruella et al. 2012). Together with Fungi and their unicellular relatives (known as Holomycota or Nucletmycea; Brown et al. 2009; Liu et al. 2009) they form the major eukaryotic supergroup Opisthokonta (Cavalier-Smith 1998; Medina et al. 2001; Adl et al. 2019). To date, five major groups of unicellular Holozoa have been recognized and characterized: Choanoflagellatea, Filasterea, Ichthyosporea, Corallochytrea (also called Pluriformea), and the recently proposed Atreyea (Torruella et al. 2015; Mitsi et al. 2026). These groups exhibit a striking diversity of morphologies and life-cycles, including a variety of transient multicellular stages (Ros-Rocher et al. 2021), such as clonal multicellular colonies, aggregative multicellularity, and multinucleate coenocytes (Marshall et al. 2008; Fairclough et al. 2010; Dayel et al. 2011; Sebé-Pedrós et al. 2013; Ros-Rocher et al. 2021; Ros-Rocher et al. 2023). Comparative genomic analyses have shown that unicellular Holozoa encode many molecular components once considered animal-specific, including genes involved in cell adhesion, cell signaling, and transcriptional regulation, suggesting that LUAA was genetically much more complex than previously thought (Sebé-Pedrós et al. 2017; Ruiz-Trillo et al. 2026). The reconstruction of LUAA relies on robust phylogenetic frameworks and broad taxonomic sampling across Holozoa. Such sampling is essential as extant Holozoa lineages represent independent evolutionary trajectories, and both genomes and life cycles vary substantially among them. Secondary gene loss, for example, can obscure accurate reconstruction, since any lineage left unsampled could hold genes or traits lost elsewhere (Albalat and Cañestro 2016; O’Malley et al. 2016).

Environmental sequencing has transformed our understanding of eukaryotic diversity by revealing lineages that remain inaccessible through other approaches. For unicellular Holozoa, environmental 18S rDNA surveys have already hinted that the current framework of five major groups is incomplete, pointing to the existence of previously unrecognized, potentially deep-branching lineages (Weber et al. 2012; Del Campo and Ruiz-Trillo 2013; Del Campo et al. 2015; Arroyo et al. 2020; Shabardina et al. 2024). However, this data remains scattered, and a systematic analysis of the diversity and environmental distribution of unicellular Holozoa is missing. Moreover, recent advances in high-throughput sequencing, including long-read metabarcoding and shotgun metagenomics, now provide unprecedented opportunities to reassess this hidden diversity. In addition, the continuous expansion of curated short-read metabarcoding databases now facilitate assessments of the environmental and biogeographical distributions of uncharacterized lineages.

Here, we re-evaluated whether the different types of available environmental sequence data reveal additional major unicellular Holozoa lineages beyond the five groups with characterized representatives, and how much deeply-branching undescribed diversity occurs within the two main established groups, Choanoflagellatea and Ichthyosporea. We then assessed the geographical and environmental distributions of all identified uncharacterized lineages across available metabarcoding samples. By combining such complementary data sources, our approach links, for the first time, phylogenetic novelty to ecological and geographical context, providing a more complete view of unicellular Holozoa diversity. Because phylogenomic reconstruction of LUAA ultimately depends on genome or transcriptome data that remain unavailable for most of this diversity, this framework provides a basis for guiding future isolation and genomic characterization.

## Results

We systematically re-evaluated unicellular Holozoa diversity by screening 18S rDNA sequences from updated reference databases together with available metagenomic and long-read metabarcoding datasets (Karst et al. 2018; Berney et al. 2022; Jamy et al. 2022; Latz et al. 2022; Sayers et al. 2025; Chuvochina et al. 2026). Candidate Holozoa environmental sequences were placed within a phylogenetic framework spanning all major Opisthokonta groups, allowing uncharacterized lineages to be identified from their phylogenetic position. We defined uncharacterized lineages as high-level phylogenetic clades without any described representative. To identify promising environments for future isolation and genomic characterization, we then examined the environmental distribution, relative abundance, and biogeography of these uncharacterized lineages by searching the EukBank short-read metabarcoding database (Berney et al. 2023) for affiliated amplicon sequence variants (ASVs) (see Methods and Text S1).

### Revealing extensive and widespread uncharacterized Holozoa diversity

The overall topology of our final maximum-likelihood (ML) phylogenetic inference of full 18S rDNA sequences is broadly consistent with current Opisthokonta systematics (Torruella et al. 2012; Mitsi et al. 2026), recovering the monophyly of major clades (Fig. 1a, Fig. S1, and Data S1-S2). As expected for an 18S rDNA-based phylogeny spanning the whole Opisthokonta diversity, several deeper relationships remained unresolved, including the monophyly of Filasterea and Ichthyosporea. We recovered nine uncharacterized lineages branching within Holozoa (Fig. 1a, Fig. S1, and Data S1-S2). Two correspond to lineages previously detected in independent surveys (*i.e.*, MAOP1 and MAOP2, Del Campo and Ruiz-Trillo 2013), while seven are newly described here. Four of these were also found in short-read metabarcoding datasets and we informally name them here (*i.e.*, HOL1, HOL2, HOL3, and HOL4). The other three are currently orphan lineages only represented by a single 18S rDNA sequence (*i.e.*, JN825684, OBEP010278239, and ORJL011316691) and we leave them unnamed. HOL4 and MAOP2 are supported as sister lineages to Choanoflagellatea, but the positions of the other seven lineages remains uncertain. HOL2 weakly affiliates with Ichthyophonida, and HOL1 and the orphan lineages with Filasterea, while MAOP1 and HOL3 have no affiliation. In addition, we found a previously unnoted and diverse uncharacterized lineage branching within Opisthokonta, but outside both Holozoa and known Holomycota clades, that we informally name OPI1 (Fig. 1a, Fig. S1, and Data S1-S2).

Based on affiliated ASVs from the EukBank database, the uncharacterized lineages are broadly distributed across both terrestrial and marine samples, indicating that they are not restricted to rare environments (Fig. 1b and Data S3-S4). However, individual lineages differ in their environmental distributions. OPI1 is primarily found in soils where it has high relative abundances, comparable to those of described unicellular Holozoa groups. On the other hand, MAOP1 seems restricted to marine environments, occurring in organism-associated, sediment, and water samples, with its highest relative abundance in the latter. MAOP2 has its highest relative abundance in land water (although the number of available samples is limited for this environmental category), and in terrestrial and marine sediments. HOL2 is predominantly distributed in marine sediments, whereas HOL4 is mainly associated with land sediments. HOL1 and HOL3 both have low relative abundances across the environments in which they are detected. No affiliated ASVs were recovered for the remaining three orphan lineages, preventing assessment of their environmental distributions. Origin of the samples where the environmental 18S rDNA sequences included in Fig. 1a were taken from is consistent with the habitats where they were detected in EukBank, and is listed in Data S1; Text S2 provides brief summaries of all the uncharacterized lineages we present here, including the types of habitats where they were found.

**Figure 1.**
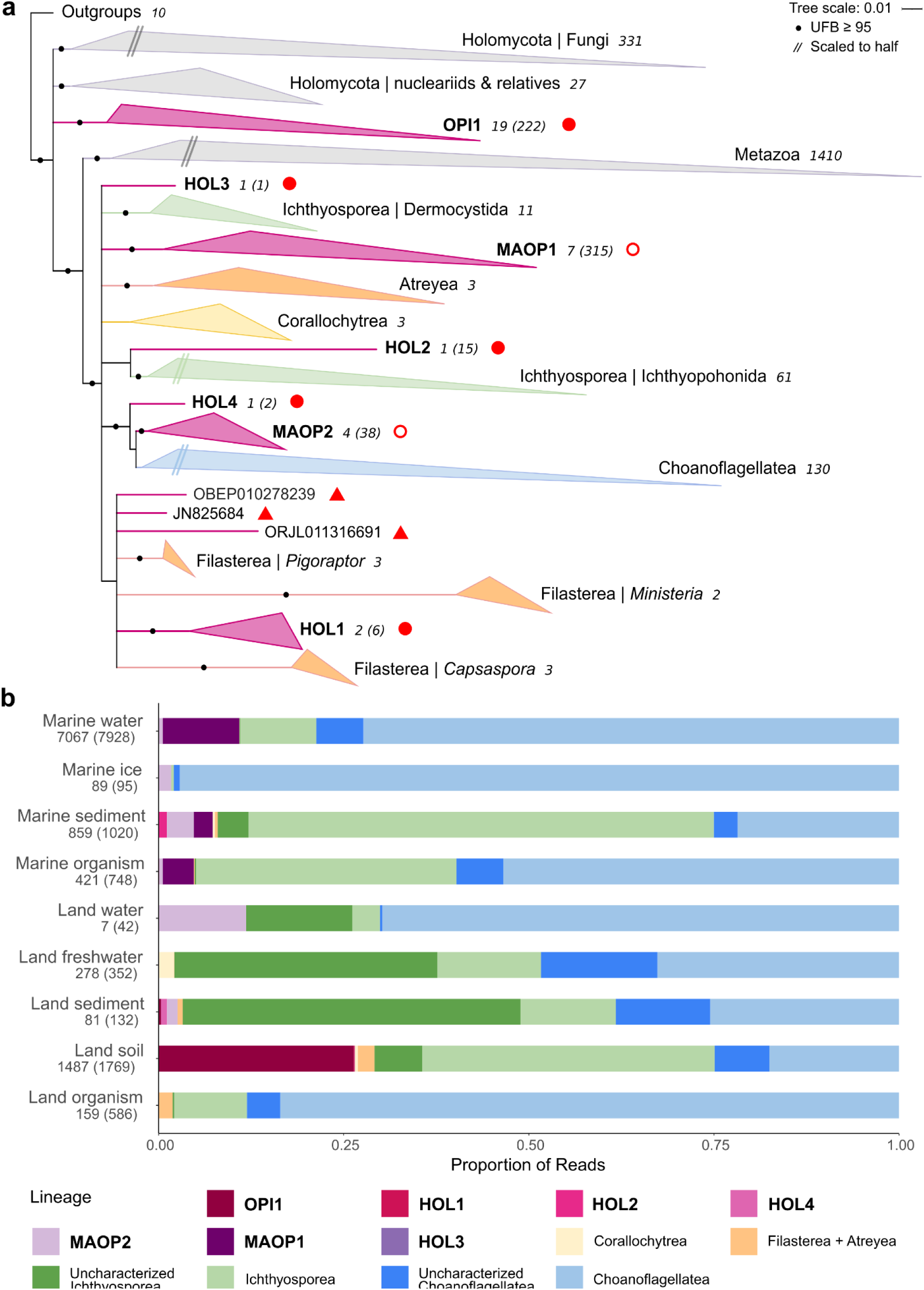
Opisthokonta phylogeny and environmental distribution of Holozoa lineages and OPI1. **a. 18S rDNA phylogeny of Opisthokonta, revealing the existence of 10 uncharacterized lineages.** The tree shown is the maximum-likelihood (ML) topology inferred from an alignment of 2,032 sequences (1,684 aligned nt positions after trimming) using the GTR+FO+I+R10 model of evolution and rooted by Breviatea, Apusomonadida and a closely related environmental lineage. Branches with <80 ultrafast bootstrap support (UFBoot) are collapsed. Lineages represented by two or more sequences are collapsed, with the number of sequences shown in italics. The ten uncharacterized lineages are shown in pink. Those identified in the EukBank database (allowing their environmental distribution to be assessed) are marked with a red circle (the second number in parentheses indicates the number of EukBank ASVs). Filled red circles denote the five lineages newly described here, whereas open red circles denote the two previously recognized lineages. Red triangles indicate the three additional, newly identified uncharacterized lineages that were not detected in EukBank and are represented by orphan sequences. The diversity inside Choanoflagellatea and Ichthyophonida is shown in Fig. 2. The uncollapsed tree with all sequence identifiers is provided in Fig. S1; see also Data S1-S2. **b. Relative abundances of EukBank ASVs affiliated to described unicellular Holozoa and uncharacterized lineages across EukBank environmental categories.** OPI1 and the six uncharacterized lineages that are putatively deep-branching within Holozoa and have affiliated ASVs are shown individually, whereas uncharacterized lineages inside Choanoflagellatea and Ichthyosporea are pooled for comparison. The “Land” categories cover all terrestrial environments, with “Land water” comprising unspecified inland aquatic environments, including saline environments. The “Marine” categories cover marine environments. For each environmental category, the number of EukBank samples where unicellular Holozoa or uncharacterized lineages were detected is indicated (with the total number of samples available in that category given in brackets). The full list of ASVs affiliated to uncharacterized lineages is provided in Data S3 and the read counts aggregated by lineage and environmental category in Data S4. **Alt text**: Figure highlighting 10 of the uncharacterized lineages we identified, with two panels a and b, providing a graphic representation of their evolutionary position within Opisthokonta (phylogenetic tree) and barplots of their relative abundances across various environments compared to known groups.

### Numerous uncharacterized lineages within Choanoflagellatea and Ichthyosporea

We next examined the environmental diversity within the two main established Holozoa groups, Ichthyosporea (which comprises two deeply diverging clades, Ichthyophonida and Dermocystida), and Choanoflagellatea. We recovered six uncharacterized lineages branching within Ichthyophonida; four (*i.e.*, FRESHIP1, FRESHIP2, SOILIP1, and MAIPh1) were found in prior environmental surveys (Del Campo and Ruiz-Trillo 2013; Shabardina et al. 2024), the other two are orphan lineages (*i.e.*, LC220088 and clone c-2682_conseq_Otu1022_5) that we leave unnamed (Fig. 2a, Fig. S1, and Data S1-S2). MAIPh1 and c-2682_conseq_Otu1022_5 are supported as sister to the clade comprising *Creolimax* and *Sphaeroforma*, FRESHIP2 is supported as sister to the CABEPI clade (comprising *Abeoforma*, *Pirum*, and *Caullerya*), FRESHIP1 is supported as sister to the AEI clade (comprising *Amoebidium*, *Ichthyophonus*, *Paramoebidium*, and Eccrinales), while SOILIP1 and LC220088 have no supported affiliations. In contrast, we did not recover additional lineages within Dermocystida.

Within Choanoflagellatea, we recovered 15 uncharacterized lineages, including six (*i.e.*, MACHO2, MACHO4, CladeL, FRESCHO2, FRESCHO3, FRESCHO4) reported in prior surveys (Weber et al. 2012; Del Campo and Ruiz-Trillo 2013), and nine newly identified. Among the latter, six are found in short-read metabarcoding datasets and informally named here (*i.e.*, CHO1, CHO2, CHO3, CHO4, CHO5 and CHO6) and three are orphan lineages (*i.e.*, GU290101, JAKFZD010052931, and JBGOPN0101245395) (Fig. 2b, Fig. S1, and Data S1-S2). Deep relationships among Choanoflagellatea are poorly resolved with 18S rDNA. We recover with maximum support the monophyly of Acanthoecida (the group comprising all species that possess a lorica) with strong support, but the diversity of species lacking alorica (classically grouped in Craspedida) is not recovered as monophyletic. Also notable is the well documented polyphyly of genus *Salpingoeca*, whose constitutive species branch in many different places across the various Craspedida lineages (Carr et al. 2017). We find that four uncharacterized lineages (FRESCHO3, CladeL, MACHO4, and MACHO2) are supported as forming a clade with Acanthoecida, and among them, MACHO4 and MACHO2 are supported as the successive sister lineages to Acanthoecida. Among the Craspedida diversity, CHO3 and CHO4 are supported as successive sister lineages to *Salpingoeca prava*, CHO2 is supported as sister to the *Mylnosiga* lineage, FRESCHO2 is supported as sister to the *Salpingoeca punica* lineage, and CHO6 is supported as sister to the *Codosiga* lineage. By contrast, CHO1, CHO5, and FRESCHO4 showed no supported affiliation. The position of the three orphan lineages is more difficult to interpret; in spite of high UFBoot support they had low transfer bootstrap expectation values (Fig. S1 and Data S2) and therefore we also regard them as lacking a supported affiliation.

**Figure 2.**
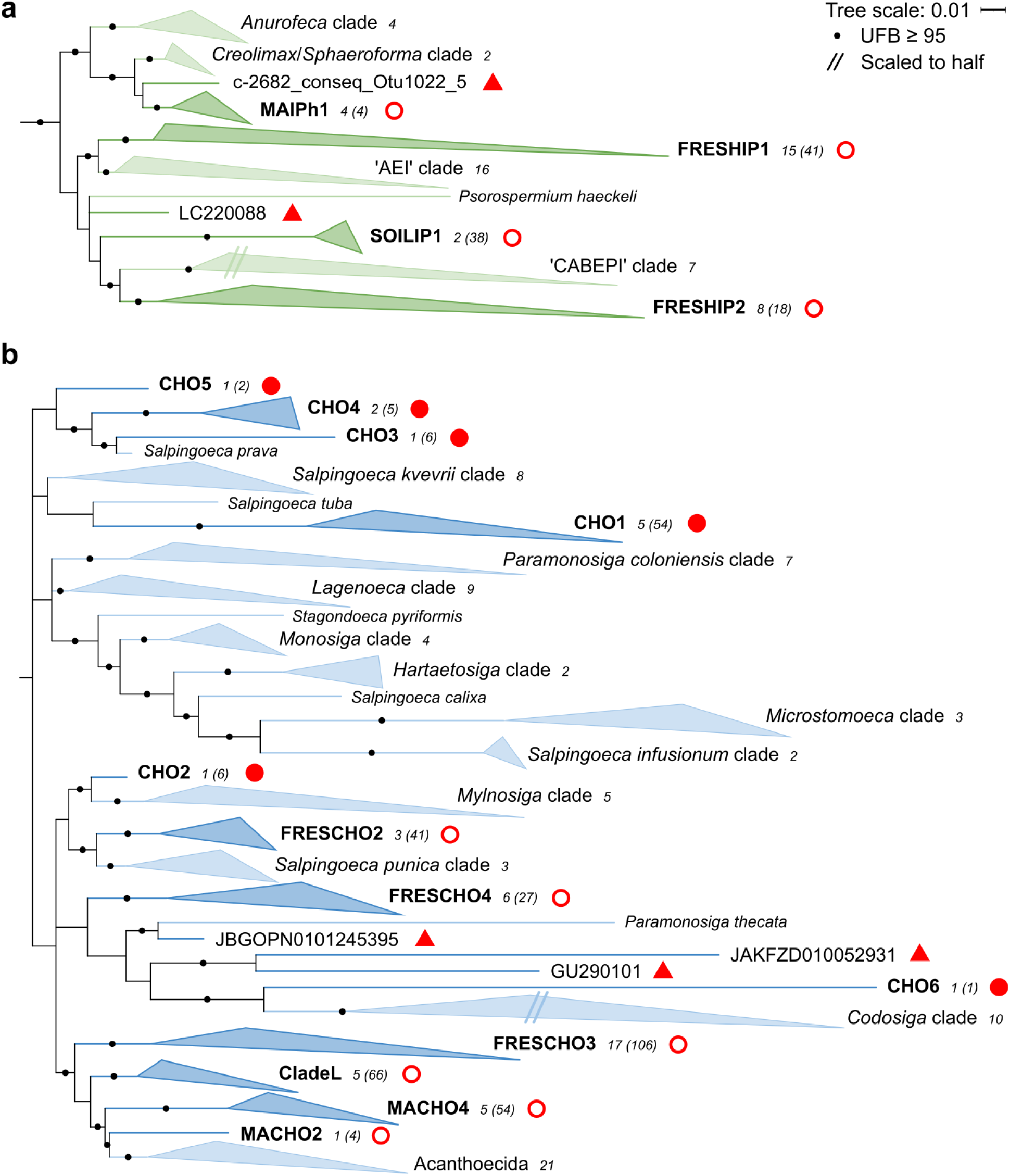
18S rDNA phylogeny of Ichthyophonida (a) and Choanoflagellatea (b), highlighting the uncharacterized lineages among them. Both panels are derived from the ML tree shown in Fig. 1a and follow the same visual conventions for collapsed branches and clades. Environmental lineages are shown in darker shades. Among the uncharacterized lineages, four in panel a and 12 in panel b were identified in the EukBank database and are marked with red circles (the second number in parentheses indicates the number of EukBank ASVs). Filled red circles denote six lineages described here for the first time, open red circles denote previously identified lineages. Red triangles indicate five additional putative uncharacterized lineages that were not detected in EukBank and are represented by orphan sequences. The uncollapsed tree with all sequence identifiers is provided in Fig. S1; see also Data S1-S2. **Alt text:** Two-panels figure providing a graphic representation (phylogenetic trees) of the diversity inside the groups Ichthyophonida and Choanoflagellatea (same analysis as in Fig. 1), highlighting the existence of six and 15 uncharacterized lineages within these groups, respectively.

Uncharacterized Ichthyosporea and Choanoflagellatea lineages accounted for a substantial fraction of reads in several environments (Fig. 1b and Data S3-S4). In particular, uncharacterized Ichthysporea matched the combined relative abundance of all known unicellular Holozoa lineages in land sediment samples, and also had higher relative abundances than characterized Ichthysporea lineages in freshwater and land water. They are also present in soil and marine sediments, but at lower relative abundances. Uncharacterized Choanoflagellatea had lower relative abundances overall, but were detected across most surveyed environments, with appreciable relative abundances in both marine and terrestrial habitats, except land water and ice.

### ASV-level analyses reveal ecological specialization in uncharacterized lineages

We then examined how ASV diversity within the uncharacterized lineages was distributed across environments. To this end, we reconstructed a ML phylogenetic tree expanded from the one shown in Fig. 1a by including all EukBank ASVs that we identified as affiliated to the uncharacterized lineages (Fig. 3, Fig. S2, and Data S3-S5). As expected, due to the addition of hundreds of short metabarcoding sequences, the phylogenetic resolution obtained with this expanded sequence dataset was lower. In particular, MAOP2 was not recovered as monophyletic. Therefore, we only use this analysis to visualize the ASV diversity within uncharacterized lineages and their distribution across environmental categories, but retain the topology of the tree shown in Fig. 1a as our reference phylogenetic framework for Holozoa relationships.

**Figure 3.**
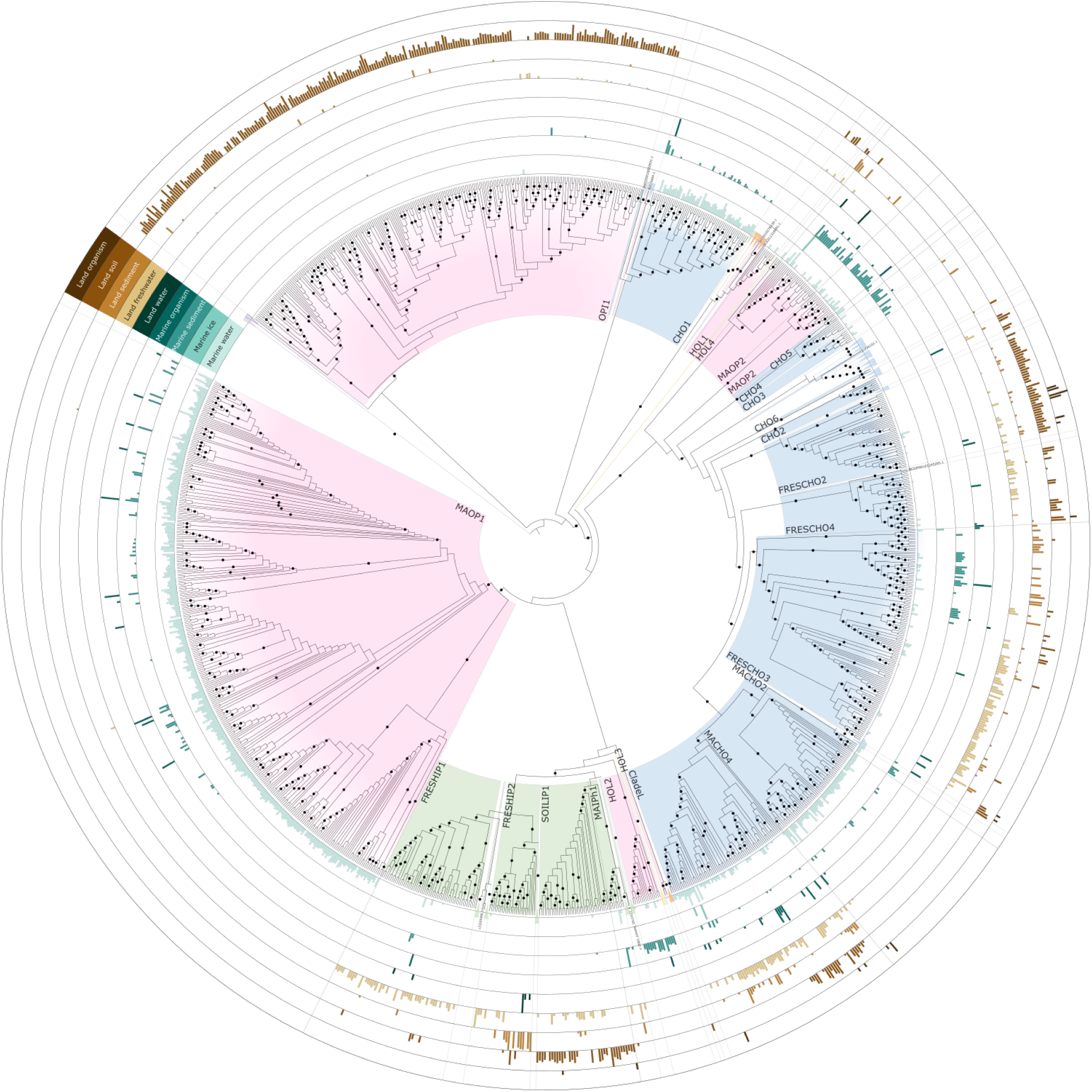
Phylogenetic diversity of the EukBank ASVs affiliated to uncharacterized lineages, and their environmental distribution across EukBank environmental categories. The tree shown is the ML topology inferred from an alignment of the 2,032 full 18S rDNA sequences present in Fig. 1a plus the 1,184 EukBank ASVs, under the GTR+FO+R10 model of sequence evolution (1,784 aligned nt positions after trimming). The tree is rooted with Breviatea, Apusomonadida, and a closely related environmental lineage. Nodes receiving ≥90 UFBoot support are indicated by black circles. Where possible, described lineages were collapsed and represented as triangles to simplify the tree. Triangle colors correspond to those used in Fig. 1a. For each EukBank environmental category and for all EukBank ASVs, the colored barplots show the log-scaled read abundance normalized to values between 0 and 1 (see Methods and Text S1). The uncollapsed tree with all sequence identifiers is provided in Fig. S2; see also Data S3-S5. **Alt text:** Circular representation of the phylogenetic tree of Opisthokonta, highlighting the ASV diversity inside all uncharacterized lineages, colored by group. For each ASV, colored bars arranged in concentric circles around the tree show their distribution across environmental categories.

Our results reveal that OPI1 is a diverse lineage within Opisthokonta based on ASVs (222), almost all of which are exclusively found in soil. In terms of the uncharacterized lineages within Holozoa, MAOP1 has the highest diversity based on ASVs (315), which are primarily restricted to marine water, although some are also found in marine sediment and marine organisms. ASVs affiliating with MAOP2 (38) and HOL2 (15) are all found in marine sediments, with some presence also in marine water. The few HOL1 ASVs (6), on the other hand, are found in distinct but diverse environments including soil, land freshwater, and marine water. We detected even lower ASV diversity for HOL3 (1) and HOL4 (2) in our survey.

Among the uncharacterized lineages inside Ichthyophonida, we found that FRESHIP1 ASVs (41) are primarily restricted to land freshwater samples. FRESHIP2 ASVs (18) fell into two main subclades, one restricted to land freshwater samples and the other found in land sediment, and also in either soil or marine organisms. SOILIP1 only comprises acouple long-read 18S rDNA sequences, but has a high diversity in terms of ASVs (38), which are primarily found in soil, with a number also detected in land freshwater. MAIPh1 ASVs (4) were detected in both marine and freshwater environments. Despite the host-association of a number of characterized Ichthyosporea lineages (Shabardina et al. 2024), very few ASVs were detected in marine or land organism samples.

Generally, uncharacterized Choanoflagellatea lineages have higher diversity at ASV level, despite being represented by only a few long-read 18S rDNA sequences in a number of cases. Several of these lineages also do not appear restricted to specific environments. For example, CladeL ASVs (66) are found in marine, freshwater, and soil environments. Similarly, ASVs affiliated with FRESCHO2 (41) are primarily found in soil, but one subclade is found in land organism samples, suggesting the potential for choanoflagellates that are host-associated. FRESCHO3 ASVs (106) have one clade primarily found in freshwater and deeper branching clades found in marine sediment or soil, while some FRESCHO4 ASVs (27) are found in freshwater and others in soil. Though with limited diversity, CHO2 ASVs (6) are primarily found in soil, but a small subclade is instead found in land freshwater. The remaining Choanoflagellatea lineages are more restricted to specific environments, with ASVs from MACHO2 (4) and CHO1 (54) found in both marine water and sediment, MACHO4 (54) restricted to marine water, CHO3 (6) to marine sediment, and CHO4 (5) to land sediment. Very limited ASV diversity was detected in our survey for CHO5 (2) and CHO6 (1).

### Global distributions identify targets for future sampling

Finally, we also used the EukBank data to examine the geographical distributions of all uncharacterized lineages by assembling for each a map displaying the environmental and geographical distributions of the affiliated ASVs alongside within-sample relative abundances (Fig. 4, Fig. S3 to S5, and Data S6). Several environmental lineages were recovered across multiple continents and oceanic regions, whereas others appeared geographically restricted. More specifically, we found that OP1 is globally distributed in soils. MAOP1, MACHO4, and CHO1 are globally distributed across the open ocean, with MAOP1 also found near the coast where it has its highest relative abundances. MAOP2 is also primarily found in coastal samples, although sometimes present in the open ocean, and in particular in Arctic samples. FRESCHO3 and CladeL are found in both land environments and marine water, but for the latter primarily in coastal samples. Based on the available data, FRESCHO2 is mainly found in European soil samples, sometimes in association with land organisms. A large number of lineages were found globally, but with a patchy presence, namely FRESHIP1, FRESHIP2, SOILIP1, MAIPh1, FRESCHO4, MACHO2, CHO2, CHO3, CHO4, and CHO5. HOL1, HOL3, and HOL4 were detected at too few locations to make inferences about their distributions and the relative abundances of CHO6 were below our cutoffs for mapping. Text S2 summarizes the diversity and environmental distribution of all uncharacterized lineages.

The relative abundances of several lineages surpassed 1% of the total eukaryotic reads in some samples, namely OPI1 (6), MAOP1 (37), CladeL (10), FRESCHO3 (2), MACHO4 (1), CHO1 (1), and FRESHIP2 (12) (Data S3). OPI1 has its highest relative abundance (3.9%) in a tropical lowlands soil sample from Sumatra, Indonesia. The highest relative abundances of FRESCHO3 (8.1%) and FRESHIP2 (9.7%) are both in freshwater lake samples, from a shallow Pampean lake in Argentina and Lake Sanabria in Spain, respectively. The highest relative abundances of MAOP1 (7.2%), CladeL (1.6%), MACHO4 (3.2%), and CHO1 (1.4%) were all found in marine water samples, from the Blanes Bay Microbial Observatory in Spain, the Baltic Sea, the South Pacific Ocean, and the Arabian Sea, respectively. With the exception of CHO1, the remaining Holozoa and Choanoflagellatea lineages newly described here had lower relative abundances, but were generally found in environment categories with fewer overall samples. The highest relative abundances of HOL1, HOL3, CHO2, and CHO6 were found in soil samples, HOL4 and CHO3 in sediment from saline springs, HOL2 from a rock sample, and CHO5 in a deep marine water sample (Data S3).

**Figure 4.**
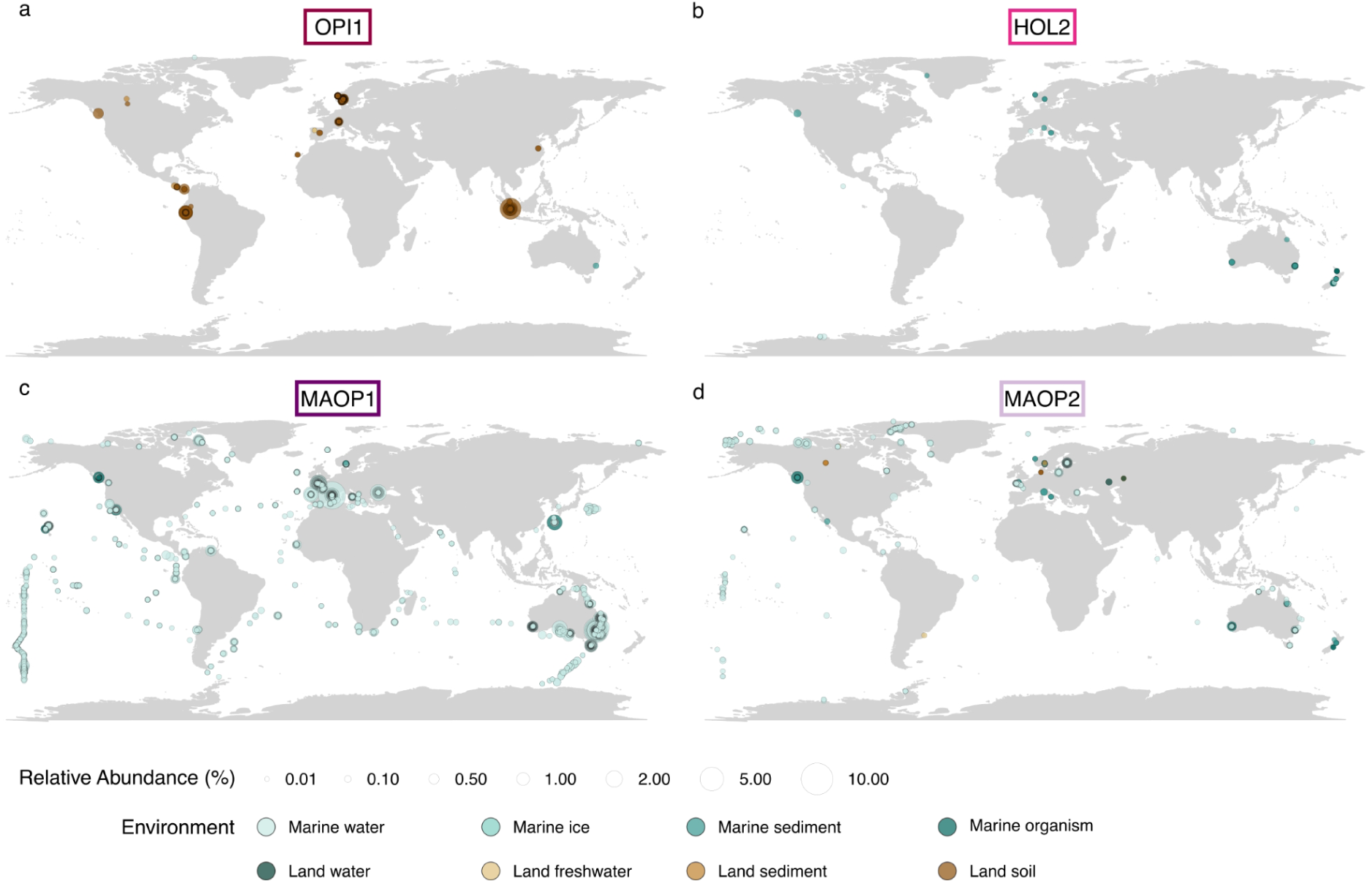
Geographical and environmental distribution of selected uncharacterized lineages. Each lineage is displayed on a separate map (a-d). Circle size is proportional to the relative abundance of reads within each sample, and circle colors indicate the EukBank environmental category. Only samples with relative abundance ≥0.01% are shown. Overlapping circles indicate multiple samples from the same location. Individual maps for the other uncultured lineages with affiliated EukBank ASVs are provided in Fig. S3 to S5; see also Data S6. **Alt text:** World maps showing the geographical location and environmental distribution of the samples containing ASVs affiliated to uncharacterized lineages OPI1, HOL2, MAOP1 and MAOP2 (panels a to d).

## Discussion

### Expansion of the diversity across Holozoa and a novel Opisthokonta lineage

Our results show that characterized unicellular Holozoa represent only a fraction of the phylogenetic diversity that can be recovered from existing environmental sequence data. Through rigorous phylogenetic validation, we here identify nine uncharacterized lineages within Holozoa that cannot be assigned to any described group, alongside an even deeper branching lineage (OPI1) at a taxonomic level comparable to Fungi and Holozoa themselves. In addition, we show the existence of 21 uncharacterized lineages within Choanoflagellatea and Ichthyosporea. Despite using a conservative approach to define uncharacterized lineages, we found 23 in multiple types of datasets, supporting their authenticity, and thereby gave them informal names. Among the uncharacterized lineages, we identified 14 of particular interest as targets (i.e., OPI1, MAOP1, MAOP2, HOL2, CladeL, FRESCHO2, FRESCHO3, FRESCHO4, MACHO4, CHO1, FRESHIP1, FRESHIP2, SOILIP1, and MAIPh1) because they (1) exhibit a high level of ASV diversity, (2) have a global distribution, and/or (3) have high relative abundances in some samples; three of these lineages are newly described here (i.e., OPI1, HOL2, CHO1). A number of these lineages have higher relative abundances than described Holozoa groups, such as Corallochytrea, Filasterea and Atreyea, and occur in various environments, highlighting that they are not rare oddities but likely represent key microbial community members whose characterization will be of ecological importance.

Much is unknown about the remaining 17 uncharacterized lineages (i.e., HOL1, HOL3, HOL4, MACHO2, CHO2, CHO3, CHO4, CHO5, CHO6, and the eight orphan lineages), all but one of which were newly identified in our study. Based on ASVs, they have limited diversity, lower relative abundances, detection in relatively few environmental samples, and in the case of the orphan lineages, were missing altogether. Results that raise the question of whether these lineages are truly rare or if they are simply poorly represented in currently available datasets. Given that these lineages were often retrieved from and detected in environments with fewer available samples, such as soil, sediments, groundwater, and deep marine water, we favour the latter explanation. Our phylogenetic analyses show that these lineages have key phylogenetic positions within Holozoa, and gathering more information about them will be crucial for gaining a more complete picture of diversity for reconstructing animal origins.

Given the wide diversity in morphologies and lifestyles across the comparatively small number of described Holozoa groups (Ros-Rocher et al. 2021), uncharacterized lineages are likely to harbor phenotypes or gene content not yet observed in Holozoa. In particular, novel insights might be expected from lineages hailing from environments with thus far little to no described species, such as sediments and soil. In this context, and due to its likely evolutionary importance, we included OPI1 in our analyses. Our phylogenetic results suggest that OPI1 is at least as distantly related to Fungi as their known relatives (the nucleariids), and may even represent a new Opisthokonta group separate from both Holozoa and Holomycota. OPI1 is widespread in soil samples, with high relative abundances, and characterization of the lifestyles and morphologies of its members is likely to be highly informative for improving our understanding of Opisthokonta evolution as a whole. While not our focus, it is also noteworthy that we captured a high number of environmental sequences phylogenetically assigned to Holozoa groups with characterized members, suggesting large untapped diversity at various taxonomic levels.

### Could additional Holozoa diversity remain undetected?

It remains to be seen how much additional Holozoa diversity we are still missing. Given our exhaustive and permissive approach, we do not expect to have missed obvious additional Holozoa lineages in the long-read 18S rDNA datasets and metagenomes we screened. However, many uncharacterized lineages were detected from undersampled environments with limited associated data. Furthermore, with the exception of surface marine water, the global distribution of the uncharacterized lineages is generally patchy. Together, this suggests that such lineages are not well captured by currently available environmental sequence datasets. Additional uncharacterized Holozoa lineages may very well remain hidden in environments with little representation. Several factors can explain these patterns. Technical biases in relation to the PCR primers used or the presence of introns or variable sequence expansions in rDNA can explain low relative abundance and prevalence, and indeed absence, of entire lineages in sequence datasets (e.g., Vaulot et al. 2022; Useros et al. 2024). And indeed it is noteworthy that five of the new uncharacterized lineages (i.e., HOL2, HOL3, CHO3, CHO4, and orphan lineage OBEP010278239) were composed solely or almost solely of long-read sequences from a PCR-free survey (Karst et al. 2018).

Beyond technical biases, the habitat representation of 18S rDNA environmental datasets remains uneven and strongly biased towards surface marine water samples. Reflecting this, the number of marine water samples in EukBank is nearly double that available for the other eight environmental categories combined. Many habitat types remain undersampled, such as deep-sea environments, coastal marine sediments, soils, freshwater sediments, or so-called “extreme” habitats (anoxic, hypersaline, acidic, etc.). Other biases are also likely at play, such as the well known issue of uneven geographical coverage, with a strong sampling bias towards temperate regions of the Northern hemisphere (Veach et al. 2021; Metz et al. 2022; Sarmento et al. 2025), which would contribute to the patchy global distributions of uncharacterized lineages. In addition, specific lifestyles, such as symbiosis or host-association, can limit the detection of lineages (Bass et al. 2015; Bass and Del Campo 2020; Burki et al. 2021). Of particular relevance to our results, is the absence of any novel lineage diversity in Dermocystida, the second subclade within Ichthyosporea, whose characterized members have a predominantly parasitic lifestyle (Rowley et al. 2013; Shabardina et al. 2024). It is likely that our search missed novel Holozoa with symbiotic or host-associated lifestyles. Lastly, we must consider the possibility that some lineages are simply genuinely rare across most or all environments. While we identified 30 uncharacterized lineages of interest within Holozoa, additional diversity most certainly remains to be uncovered.

### From detection to characterization of the uncovered Holozoa diversity

The identification of deep uncharacterized diversity within Holozoa and of numerous uncharacterized lineages inside Choanoflagellatea and Ichthyophonida is only a first step towards a greater goal of characterizing the full breadth of phenotypic and genomic diversity that exists among the close relatives of animals. The next step is to isolate and describe representatives of these uncharacterized lineages. By linking for the first time new phylogenetic diversity to geographical and environmental distributions, our study provides a clear roadmap for targeted sampling and isolation efforts.

Due to their large diversity, global distribution, and/or high relative abundance, the 14 lineages highlighted above represent prime targets for such isolation efforts. However, therein lies a paradox: the fact that such lineages have so far remained uncharacterized suggests they may prove difficult to isolate and culture. Nevertheless, a number of these lineages have high enough relative abundances that in such cases, partial morphological characterization should remain achievable through fluorescent in-situ hybridization (Massana et al. 2002; Massana et al. 2004; Not et al. 2007), and genomic characterization through cell sorting approaches coupled with single-cell transcriptomics or genomics (Heywood et al. 2011; Krabberød et al. 2017; Mangot et al. 2017). Metagenome resolved genomics may even be an option in samples with high relative abundances of the target lineage (Delmont et al. 2022; Alexander et al. 2023), as was recently applied to obtain the genome of the parasite *Txikispora philomaios* and to thereby delineate the Holozoa clade Atreyea (Mitsi et al. 2026).

### Towards reconstructing the phenotype and genotype of LUAA

Our study doubles the number of known deep lineages that are close relatives of animals, bringing us to nine Holozoa lineages lacking genomic and morphological characterization (i.e, MAOP1, MAOP2, HOL1, HOL2, HOL3, HOL4, and the three orphan lineages), and missing from current comparative studies of animal origins. Yet, broad taxon sampling is key for accurately reconstructing the phenotype and genotype of LUAA (Suga et al. 2012; Richter et al. 2018; Ocaña-Pallarès et al. 2022; Ruiz-Trillo et al. 2023). Efforts targeting the isolation and characterization of these lineages are now crucial, and will likely lead to novel insights into the origin of animals. In addition, given the broad phenotypic, genomic, and life cycle diversity inside the established Holozoa groups (Ros-Rocher et al. 2021), expanding representation within those is also of large importance.

Our phylogenetic framework to place the uncharacterized Holozoa lineages relies solely on 18S rDNA sequences, thus their exact phylogenetic positions and level of taxonomic novelty remain uncertain. As illustrated in our analyses, 18S rDNA alone fails to recover well established relationships within Holozoa, such as the monophyly of Ichthyosporea (with Dermocystida and Ichthyophonida branching separately), and typically only recovers Corallochytrea, and Filasterea each as monophyletic without statistical support (Del Campo and Ruiz-Trillo 2013; Urrutia et al. 2022; Shabardina et al. 2024). It is therefore possible that some of the uncharacterized lineages actually belong to one of the established groups, in particular Filasterea, with which several lineages affiliate. With this phylogenetic uncertainty we also cannot currently exclude that one or more of these lineages may prove to represent a closer sister-group to Metazoa than Choanoflagellatea. The tantalizing possibility further highlights the crucial importance of genomic characterization of the full lineage diversity within Holozoa.

Phylogenomics remains the most powerful approach for resolving deep evolutionary relationships and reconstructing ancestral states, but its power is inherently bounded by the diversity it can draw on: a phylogenomic and comparative genomic reconstruction of LUAA can only be as complete as the sampling that feeds it. By mapping where the missing diversity lives, our results turn a previously untraceable gap into a concrete set of targets. The key to understanding the unicellular origins of animal multicellularity may already be within reach: surrounding us, still largely unknown.

## Methods

The detailed methodology we followed is described in Text S1 and summarized below.

### Searches for uncharacterized lineages in environmental datasets

Eukayotic 18S rDNA reference sequences were retrieved from EukRibo v2.0 (Berney et al. 2022), with the addition of the *Txikispora philomaios* 18S rDNA (NCBI accession: OK181898; Urrutia et al. 2022), and used later in phylogenetic analyses, and the subset of Holozoa sequences for dataset searches (Text S1). The region corresponding to the 18S rDNA was extracted from EukRibo sequences using Barrnap v0.9 (Seemann 2018) with the option “--kingdom euk”. Different identity thresholds were used to cluster 18S rDNA sequences from various taxonomic groups to reduce sequence redundancy using CD-HIT-EST v4.8.1 (Li and Godzik 2006; Fu et al. 2012). Potential environmental Holozoa diversity was then retrieved through blastn v2.12.0+ (Altschul et al. 1990; Camacho et al. 2009) searches of the Holozoa reference sequences, with ≥80% sequence identify and query coverage of ≥30%, against the following databases: the NCBI core nucleotide BLAST database (core_nt, version 5, as of January 2025; (Sayers et al. 2025)), NCBI metagenomes (as of March 2025, Sayers et al. 2025), the SILVA SSU Parc database (taxonomy release 138.2; (Chuvochina et al. 2026)), a dataset of PCR-free environmental rDNA sequences (Karst et al. 2018), and two long-read environmental rDNA datasets generated with universal eukaryotic primers (Jamy et al. 2022; Latz et al. 2022) (Text S1). The long-read data from Latz et al. 2022 was first processed from raw sequencing reads using the pipeline outlined in Jamy et al. 2022, except for filtering of reads between 2,000 and 6,000 bp prior to OTU clustering. Sequences from the Karst et al. 2018 dataset were first clustered at 99% identity to remove redundancy using CD-HIT-EST v4.8.1 (Li and Godzik 2006; Fu et al. 2012). The 18S rDNA region was then extracted from all non-redundant sequence hits as previously, and resulting sequences shorter than 700 bp discarded with SeqKit v2.8.2 (Shen et al. 2024). Remaining sequences were then clustered at 97% identity with VSEARCH v2.29.1 (Rognes et al. 2016) with the options --cluster_fast and --centroids, resulting in 23,869 18S rDNA sequence clusters from which to identify novel environmental Holozoa diversity. Blastn v2.12.0+ was then used to eliminate sequences outside of the taxonomic scope of the study. Those with a top hit to EukRibo (Berney et al. 2022) database sequences outside Opisthokonta or within Fungi or Bilateria with ≥95% sequence identity and ≥80% coverage were removed, resulting in 12,262 18S rDNA sequences for phylogenetic inspection.

### Phylogenetic identification of uncharacterized Holozoa lineages

A multi-step phylogenetic filtering approach was then used to identify uncharacterized Holozoa, or Opisthokonta, lineages (Text S1) - first by inferring iterative eukaryote-wide phylogenies with a subset of EukRibo v2.0 (Berney et al. 2022) reference sequences to further exclude non-target 18S rDNA sequences, then second by additional iterations with a subset of EukRibo Opisthokonta references and select outgroups. In all cases, EukRibo reference sequences were aligned using MAFFT v7.525 (Katoh and Standley 2013) with the “--maxiterate 1000 --auto” options, and candidate sequences then added to the reference alignment with MAFFT’s “--add” option. TrimAl v1.5.rev0 (Capella-Gutiérrez et al. 2009) was used to trim alignments with the “-gt 0.5” option (Text S1). ML phylogenetic trees were then inferred using IQ-TREE v2.3.6 (Minh et al. 2020) with model selection using ModelFinder (Kalyaanamoorthy et al. 2017) (“-m MFP”) from among GTR (Lanave et al. 1984; Tavaré 1986) base substitution models (“-mset GTR”), with empirical or ML optimized base frequencies (“-mfreq,F,FO”)”, and 1000 ultra-fast bootstrap replicates (Hoang et al. 2018) (“-B 1000”). Prior to inference of the Opisthokonta phylogenetic trees, 1,228 candidate sequences retained after the eukaryote-wide filtering step were also subjected to two independent chimera checks, using the UCHIME algorithm v4.2 (Edgar et al. 2011) from Mothur v1.39.5 (Schloss et al. 2009) (67 putative chimeras) and VSEARCH (in reference mode) with the SILVA SSU Ref NR99 database (235 putative chimeras and 506 borderline sequences). Only sequences classified as non-chimeric and not borderline by both methods were retained, resulting in 1,016 candidate Holozoa 18S rDNA sequence clusters whose phylogenetic positions in Opisthokonta were then examined. Through iterative phylogenies, we refined our candidate selection, retaining only sequences representing completely uncharacterized diversity (*i.e.*, forming lineages composed entirely of environmental sequences) branching within Opisthokonta, Holozoa, or deeply within one of the unicellular Holozoa lineages. Our phylogenetic filtering efforts resulted in the identification of the 31 uncharacterized lineages we present in this study, which are composed of 120 18S rDNA sequence clusters (corresponding to 250 individual sequences; see Data S1). For the final phylogenetic inference, we also retained 40 sequence clusters branching inside characterized lineages with limited described diversity, such as Corallochytrea, Filasterea, and Atreyea (see Text S1). Branch support was then also evaluated using the transfer bootstrap expectation (TBE; (Lemoine et al. 2018)) estimation after 300 bootstrap replicates (-b 300 -tbe -undo). The resulting TBE values (shown in Fig. S1; see also Data S2) were mapped onto the final ML topology using RAxML-NG v1.2.2 (Kozlov et al. 2019) (--support --bs-metric tbe).

### Geographical and environmental distributions of uncharacterized lineages

The EukBank v4 18S rDNA short-read metabarcoding database (Berney et al. 2023) was screened to identify ASVs affiliated with the 31 uncharacterized lineages. Representative sequences from each of the 120 clusters, corresponding to the uncharacterized lineages, were used to search EukBank with Blastn v2.12.0+ (Altschul et al. 1990; Camacho et al. 2009) and ASVs with ≥85% and ≥5% query coverage retained, resulting in 12,272 EukBank ASVs. We then applied the same iterative filtering steps used previously (reverse BLAST and phylogenetic verification) to exclude ASVs not affiliated to the 31 uncharacterized lineages, except ASVs were added to reference alignments using MAFFT’s “--addfragments” option (Text S1). EukBank ASVs are connected to hierarchical single linkage clusters based on pairwise distances, which were used to add ASVs linked to those identified in the BLAST searches and phylogenetically verified (Text S1). ASVs were verified as affiliated with one of the 31 uncharacterized lineages if they branched together with ultrafast bootstrap support ≥90 to the exclusion of any EukRibo reference sequence. The iterative ML phylogenetic filtering validation steps resulted in a final selection of 1,072 ASVs affiliated to one of the 31 uncharacterized lineages (Data S3). Relative abundances of characterized unicellular Holozoa and uncharacterized lineages across environments were estimated using EukBank metadata. For characterized unicellular Holozoa, ASVs were selected using the “taxogroup1” assignments, while the previously identified affiliated ASVs were used for the uncharacterized lineages. Read counts for each ASV were then summed across samples and environmental categories in which they were present (Text S1). When an ASV affiliated to an uncharacterized lineage was also classified as a described Holozoa group, reads were counted only toward the uncharacterized lineage. The geographic distribution of the uncharacterized lineages was also examined using EukBank metadata. For each lineage, the coordinates of samples including read counts from affiliated ASVs were retrieved. Relative abundances of uncharacterized lineages in each sample were calculated by dividing the number of reads from affiliated ASVs by the total sample read number. The environmental distribution of individual affiliated ASVs was calculated by summing read counts across samples for each environmental category. Read counts were then log10(x+1)-transformed and min-max scaled to values between 0 and 1 within each environmental category, across ASVs.

### Data visualization

Alignments were visualized with AliView v1.28 (Larsson 2014), phylogenetic trees with Dendroscope v3.8.10 (Huson and Scornavacca 2012), FigTree v1.4.4 (Rambaut et al. 2018), or iTOL v7.6 (Letunic and Bork 2024). R v4.5.1 (R Core Team 2025) and RStudio v2025.05.1+513 (Posit Team) were used for generating plots and maps, with the following packages: maps v3.4.3, ggplot2 v3.5.2, dplyr v1.1.4, patchwork v1.3.2, gridExtra v2.3, scales v1.4.0, data.table v1.17.8, and ggforce v0.5.0. Figures were edited in Inkscape v1.4.2 (Inkscape Project 2025). See also Data and Code Availability.

## Data and Code Availability

All data supporting the findings of this study is provided in the supplementary data files (Data S1-S6). Custom scripts for processing EukBank data are publicly available on GitHub at https://github.com/alijimv/Exploring-Unicellular-Holozoa-Diversity.

## Material and Resource Availability

All materials and resources necessary to reproduce the study findings are provided in the article and supplementary materials. Additional information and requests for resources should be directed to Iñaki Ruiz-Trillo or Cédric Berney.

## Acknowledgments

This work is supported by an ERC grant (MISSINGRELATIVES, 101097659) funded by the European Union. Views and opinions expressed are however those of the author(s) only and do not necessarily reflect those of the European Union or the European Research Council Executive Agency. Neither the European Union nor the granting authority can be held responsible for them. In addition, J.E.D. is supported by an international postdoctoral grant from the Swedish Research Council (VR grant 2022-06250) and is a non-stipendiary EMBO Postdoctoral Fellowship recipient (ALTF 740-2022). Computational analyses were enabled by resources from projects NAISS 2023/23-512, NAISS 2023/22-491, NAISS 2024/23-629, NAISS 2024/22-1422, and UPPMAX 2025/2-186 provided by the National Academic Infrastructure for Supercomputing in Sweden (NAISS) at UPPMAX, funded by the Swedish Research Council through grant agreement no. 2022-06725. We thank the Galician Supercomputing Center (CESGA) for granting access to its supercomputing infrastructure. The supercomputer FinisTerrae III and its permanent data storage system have been funded by the NextGeneration EU 2021 Recovery, Transformation and Resilience Plan, ICT2021-006904, and also from the Pluriregional Operational Programme of Spain 2014-2020 of the European Regional Development Fund (ERDF), ICTS-2019-02-CESGA-3, and from the State Programme for the Promotion of Scientific and Technical Research of Excellence of the State Plan for Scientific and Technical Research and Innovation 2013-2016 State subprogramme for scientific and technical infrastructures and equipment of ERDF, CESG15-DE-3114 We thank Miguel Méndez Sandín for advice and helpful discussions.

## Author Contributions

Conceptualization: IRT, JED, and CB. Formal analysis: AJV, JED, and CB. Funding acquisition: IRT. Investigation: AJV, JED, and CB. Methodology: AJV, JED, and CB. Resources: IRT and JED. Supervision: IRT, JED, and CB. Visualization: AJV. Writing – original draft: AJV. Writing – review & editing: AJV, IRT, JED, and CB.

