## Supplementary Information for "Uncharacterized unicellular relatives reveal key gaps in sampling diversity for reconstructing animal origins"

This document contains the following:

**Text S1: Supplementary Methods**

**Text S2: Descriptions of the Uncharacterized Lineages**

**Supplementary Figure Legends**

**Supplementary Data Descriptions**

#### Text S1: Supplementary Methods

This text details the full procedure that we followed to identify new Holozoa diversity, define uncharacterized lineages of interest by phylogenetic analyses, and explore the geographical and environmental diversity of these uncharacterized lineages using short-read metabarcoding datasets. All citations are included in the main text.

##### *Building sets of reference eukaryotic 18S rDNA sequences*

The reference 18S rDNA sequences used in this study were taken from EukRibo version 2.0 (Berney et al. 2022), a curated reference database of eukaryotic 18S rDNA sequences. EukRibo provides a fasta file of the full sequences as retrieved from their NCBI accession numbers, which can include more than just the 18S rDNA. Therefore for all of them we extracted the part corresponding to the 18S rDNA with Barrnap v0.9 (Seemann 2013), using the option "--kingdom euk". Sequences for which Barrnap yielded multiple 18S rDNA fragments (typically because of the presence of introns) were discarded, after we double-checked that they did not represent a loss of important diversity. Sequences flagged as chimeric in EukRibo were also excluded. The 18S rDNA sequence of *Txikispora philomaios* (NCBI accession number OK181898; Urrutia et al. 2022) was added to the remaining EukRibo sequences because it was published after the release of EukRibo version 2.0 (Berney et al. 2022). This yielded a starting set of 45,013 reference 18S rDNA sequences (RefSet0) from which we built four subsets of sequences used at different subsequent stages of our work.

First, we selected all sequences from unicellular Holozoa and non-bilaterian Metazoa, except Myxozoa, a parasitic cnidarian lineage with highly divergent 18S rDNA sequences. Then we clustered these sequences at 97% identity with CD-HIT-EST v4.8.1 (-c 0.97; (Li and Godzik 2006; Fu et al. 2012)). This resulted in a subset of 365 unicellular Holozoa and non-bilaterian Metazoa sequences (RefSet1) that we used as BLAST queries to identify potential novel Holozoa diversity in various sequence databases (see below). Next, we created an exhaustive, eukaryote-wide subset of sequences consisting of the full RefSet0 filtered only to exclude sequences shorter than 700 bp. This resulted in a subset of 44,875 eukaryotic sequences (RefSet2) that we used as a reverse BLAST database to exclude environmental clones that clearly belong to non-Holozoa lineages.


2

Another, more reduced eukaryote-wide subset of reference sequences (RefSet3) was then built for phylogenetic reconstructions. Several filtering steps were applied to reduce taxonomic overrepresentation while maintaining sufficient coverage of all known eukaryotic groups: (1) groups with highly divergent 18S rDNA sequences (such as Foraminifera and Euglenozoa) were excluded; (2) sequences shorter than 1,500 bp were removed; and (3) the remaining sequences were clustered with VSEARCH v2.29.1 (Rognes et al. 2016) at different identity thresholds for different taxonomic groups (97% for all Opisthokonta except Fungi and Bilateria, 95% for Fungi and non-opisthokont taxa, and 90% for Bilateria). This produced a pre-selection of 5,307 representative eukaryotic sequences. These sequences were aligned with MAFFT v7.525 (Katoh and Standley 2013) using the "—maxiterate 1000 –auto" options, and trimmed with trimAl v1.5.rev0 (Capella-Gutiérrez et al. 2009) with the "-gt 0.5" flag, retaining 1,673 aligned nucleotide positions. A maximum likelihood (ML) phylogeny was then inferred using IQ-TREE v2.3.6 (Minh et al. 2020) with the options "-m MFP -mset GTR -mfreq ,F,FO" and with 1000 ultra-fast bootstrap (Hoang et al. 2018) replicates with flag "-B 1000". The best-fit substitution model was selected from among GTR models (Lanave et al. 1984; Tavaré 1986) using ModelFinder (Kalyaanamoorthy et al. 2017), and the selected model was GTR+FO+I+R10. The resulting phylogenetic tree was arbitrarily rooted with the Discoba genus *Andalucina*. Root-to-tip distances were then calculated, and the 10% most divergent sequences (530 sequences) were removed to limit long-branch artefacts. This yielded a subset of 4,777 eukaryotic sequences (RefSet3), which were re-aligned with MAFFT v.7.525 (Katoh and Standley 2013) to improve alignment quality using the options "–maxiterate 1000 –auto –op 5 –ep 0", and then used for phylogenetic filtering of putative Holozoa environmental clones (see below).

Finally, we generated a subset of sequences restricted to Opisthokonta and closely related outgroups (Apusomonadida, Breviatea and an environmental lineage). Again differential clustering was performed with VSEARCH v2.29.1 (Rognes et al. 2016) (--cluster_fast --centroids): Holozoa (excluding Bilateria) and Holomycota (excluding Fungi) were clustered at 99% sequence identity, Fungi were clustered at 95% sequence identity, and Bilateria and outgroups were clustered at 90% sequence identity. Clusters composed of a single sequence were discarded from Fungi, Bilateria and outgroups to limit the influence of divergent sequences on phylogenetic inference. This resulted in a subset of 2,080 sequences, which were aligned and trimmed as described above, resulting in 1,741 aligned nucleotide positions. A ML phylogeny was inferred using IQ-TREE v2.3.6 (Minh et al. 2020) with the same options and evolutionary model as before. The phylogenetic tree was rooted between Opisthokonta and the outgroups. Root-to-tip distances were


3

then calculated, and the 10% most divergent sequences (208 sequences) were removed to limit the risk of long-branch artefacts. This yielded a final subset of 1,872 sequences (RefSet4), which were re-aligned with MAFFT v7.525 (Katoh and Standley 2013) as described above to improve alignment quality, and then used to build a final reference alignment to place the new Holozoa diversity in a phylogenetic context and build the tree shown in Fig. 1a (see below).

***Identification of potential novel Holozoa environmental diversity by BLAST searches***

The following public resources were screened by BLAST (Altschul et al. 1990; Camacho et al. 2009) to retrieve potential environmental Holozoa diversity: the NCBI core nucleotide BLAST database (core_nt, version 5, as of January 2025; (Sayers et al. 2025)), NCBI metagenomes (as of March 2025, (Sayers et al. 2025)), the SILVA SSU Parc database (taxonomy release 138.2; Chuvochina et al. 2026), a dataset of PCR-free environmental rDNA sequences (Karst et al. 2018), and two long-read environmental rDNA datasets generated with universal eukaryotic primers (Jamy et al. 2022; Latz et al. 2022). The 18S rDNA OTUs from Jamy et al. 2022 were retrieved from the publication's data repository (https://doi.org/10.6084/m9.figshare.15164772), whereas for the data from Latz et al. 2022 we processed the raw sequencing reads *de novo* following the pipeline used by Jamy et al. 2022, except that sequences were filtered to retain only reads between 2,000 and 6,000 bp prior to OTU clustering. A separate local BLAST search database was generated for each of these data sources using the "makeblastdb" tool.

BLAST searches were performed with blastn v2.12.0+ (Altschul et al. 1990; Camacho et al. 2009) against all data sources using as queries all the sequences in RefSet1, requiring a minimum sequence identity of 80% (-perc_identity 80), and retrieving up to 5,000 hits per query (-max_target_seqs 5000). We chose this low similarity threshold to minimize the risk of missing interesting Holozoa diversity because of 18S rDNA sequence divergence. Only matches with at least 30% query coverage and sequence lengths ≥700 bp were retained. NCBI BLAST records annotated as genome assemblies or chromosomes were excluded, as they only included organisms for which we already had a reference sequence. Redundant sequences shared across EukRibo, SILVA SSU Parc, and the NCBI core_nt databases were identified from their accession numbers. Sequences already present in EukRibo were ignored and for duplicates shared only between the other two, the SILVA record was kept. For the dataset from Karst et al. 2018, redundant sequences were collapsed at 99% identity with CD-HIT-EST v4.8.1 (Li and

Godzik 2006; Fu et al. 2012). The 18S rDNA region was extracted from all retained sequences with Barrnap v0.9 (Seemann 2013) (--kingdom euk), and sequences shorter than 700 bp were discarded using the "seq" tool from SeqKit v2.8.2 (Shen et al. 2024) with the "-m 700" option. The remaining sequences were clustered together at 97% identity with VSEARCH v2.29.1 (Rognes et al. 2016) using the options --cluster_fast and --centroids to generate a non-redundant dataset of 23,869 sequence clusters representing potential new Holozoa diversity.

***Phylogenetic filtering of the candidate clusters and construction of the reference Opisthokonta phylogeny shown in Fig. 1a***

To exclude environmental sequences outside the taxonomic scope of this study, we followed an iterative procedure relying on the other subsets of reference sequences (RefSet2, RefSet3, and RefSet4). First, the representative sequence of each of the 23,869 candidate clusters identified at the previous step were searched by reverse BLAST against our exhaustive RefSet2 using blastn v2.12.0+ (Altschul et al. 1990; Camacho et al. 2009). Sequences whose best BLAST hit (at ≥95% sequence identity and with ≥80% coverage) was outside Opisthokonta or within Fungi or Bilateria were discarded. After this step, a total of 12,262 candidate Holozoa sequence clusters were retained. The representative sequences of these 12,262 clusters were then added to the RefSet3 alignment using the MAFFT v7.525 (Katoh and Standley 2013) --add option to preserve the reference alignment, and trimming was done as before, resulting in 1,564 aligned nucleotide positions. A ML phylogeny was generated from this trimmed alignment of 17,039 sequences using IQ-TREE v2.3.6 (Minh et al. 2020) with the same options and evolutionary model as before. The tree was inspected manually in order to further eliminate all candidate clusters whose representative sequence branched outside of Opisthokonta or within Fungi or Bilateria with ultra-fast bootstrap support values ≥95%. After this first phylogenetic selection, a total of 1,228 candidate Holozoa sequence clusters were retained.

At this stage we expanded the 1,228 clusters to recover all constituent sequences and performed two independent chimera checks with Mothur v1.39.5 (Schloss et al. 2009) using the UCHIME algorithm v4.2 (Edgar et al. 2011), as well as VSEARCH v2.29.1 (Rognes et al. 2016) in reference mode, using the SILVA SSU Ref NR99 database (taxonomy release 138.2; Chuvochina et al. 2026) as the reference. Mothur identified 67 putative chimeras, whereas VSEARCH detected 235 chimeras and classified an additional 506 sequences as borderline. Conservatively, we decided to retain only sequences classified as non-chimeric by both methods. The resulting dataset comprised 9,306 sequences, which were re-clustered at 97% sequence identity with VSEARCH v2.29.1 (Rognes et al. 2016) (--cluster_fast --centroids), yielding 1,074 candidate Holozoa sequence clusters. We then repeated the phylogenetic check described above. The 1,074 clusters were re-incorporated into the RefSet3 alignment using the MAFFT v7.525 (Katoh and Standley 2013) --add option. The resulting alignment of 5,851 sequences was trimmed as before, keeping 1,684 aligned nucleotide positions, and a ML phylogeny was generated with IQ-TREE v2.3.6 (Kalyaanamoorthy et al. 2017; Hoang et al. 2018; Minh et al. 2020) as described before. Again we inspected the resulting tree manually and the same selection criteria was applied: we retained all sequences branching within Opisthokonta but outside of Fungi or Bilateria (with ultra-fast bootstrap support values ≥95%). This resulted in a final set of 1,016 candidate Holozoa sequence clusters.

The precise affiliation of these clusters within Opisthokonta was then assessed using our RefSet4, in order to determine which ones represented new diversity within lineages containing characterized organisms, and which ones represented the kind of deep novel diversity that we were interested in identifying. The representative sequences of the final set of 1,016 candidate Holozoa sequence clusters from the previous selection step were incorporated into the RefSet4 alignment using the MAFFT v7.525 (Katoh and Standley 2013) --add option to preserve the reference alignment, and trimming was done as before, resulting in 1,719 aligned nucleotide positions. A ML phylogeny was generated with IQ-TREE v2.3.6 (Minh et al. 2020) as described before, and the resulting tree was manually inspected after collapsing branches with ultra-fast bootstrap support values <90%.

Within Holomycota, sequences were retained only if they branched outside of the clades containing characterized organisms. We identified a single such lineage composed entirely of environmental sequences (uncharacterized lineage OPI1). Within Holozoa, we extended the sequence selection. We retained all the sequences branching outside of the known clades (which fell into nine uncharacterized lineages). But for Choanoflagellatea, Ichthyosporea, and non-bilaterian Metazoa, we also retained all sequences forming lineages without any reference sequence. We identified 15 such lineages in Choanoflagellatea, and six such lineages in Ichthyosporea. In non-bilaterian Metazoa, we didn’t identify any new lineage outside of the known phyla (Porifera, Ctenophora, Cnidaria, and Placozoa). This selection resulted in a total of 160 sequence clusters whose representative sequences are included in Fig. 1; of these 120 clusters (corresponding to 250 individual sequences; see Data_S1) belong to the 31 uncharacterized lineages we present here. For the final phylogenetic inference, we also retained 40 sequence clusters branching inside characterized lineages because additional sequence diversity would help with the final reference tree topology.

This final set of sequences was incorporated again into the RefSet4 alignment using the MAFFT v7.525 (Katoh and Standley 2013) --add option to preserve the reference alignment, and trimming was done as before, resulting in 1,684 aligned nucleotide positions. A ML phylogeny was inferred from this final dataset/alignment of 2,032 sequences with IQ-TREE v2.3.6 (Minh et al. 2020) as described before. To evaluate branch support beyond the 1000 ultra-fast bootstrap replicates, an additional IQ-TREE analysis was performed using 300 bootstrap replicates and a transfer bootstrap expectation (TBE; Lemoine et al. 2018) estimation (-b 300 -–tbe -undo). The resulting TBE values (shown in Fig. S1; see also Data S1) were mapped onto the final ML topology using RAxML-NG v1.2.2 (Kozlov et al. 2019) (--support --bs-metric tbe). The maximum-likelihood phylogenetic tree shown in Figs. 1a and 2, and Fig. S1 (Newick format) and the alignment used to generate it (before and after trimming, FASTA format) are provided in Data S2.

***Retrieval of EukBank ASVs associated with our uncharacterized lineages***

To obtain further information on the environmental distribution, relative abundance, and biogeography of these uncharacterized lineages and identify promising environments for future isolation and genomic characterization, we then searched the EukBank v1.0 short-read metabarcoding database, compiling sequence datasets of the variable region V4 of the 18S rDNA gene (Berney et al. 2023). To retrieve all ASVs affiliated to our uncharacterized lineages, a blastn v2.12.0+ search (Altschul et al. 1990; Camacho et al. 2009) was performed against EukBank. We used as queries the representative sequences of the 160 sequence clusters, with a minimum sequence identity threshold of 85% (-perc_identity 85) and retaining up to 1,000 hits per query (-max_target_seqs 1000). Only ASVs with ≥5% query coverage were retained for downstream analyses. This search retrieved 12,272 EukBank ASVs.

As we did before for all environmental clones, we then used several filtering steps to eliminate ASVs not affiliated to our uncharacterized lineages. First, the 12,272 ASVs were searched by reverse BLAST against RefSet2 using blastn v2.12.0+ (Camacho et al. 2009). Only ASVs whose 500 best BLAST hits (-max_target_seqs 500) did not include any reference sequence with ≥95% sequence identity were retained, which left 7,619 ASVs. The next filtering step was phylogenetic, using our final dataset/alignment of 2,032 sequences. The 7,619 ASVs were added to this alignment as fragments (MAFFT v7.525 (Katoh and Standley 2013) option --addfragments) to preserve the reference alignment. The resulting alignment of 9,651 sequences consisted of 1,843 aligned nucleotide positions after trimming with trimAl v1.5.rev0 (Capella-Gutiérrez et al. 2009) (-gt 0.1). A ML phylogeny was inferred with IQ-TREE v2.3.6 (Minh et al. 2020) as described before and the resulting tree was manually inspected. As an initial selection, all ASVs branching more closely with our query clusters than with any characterized taxon were retained, irrespective of UFBoot support values. This resulted in a selection of 1,651 ASVs potentially affiliated to uncharacterized lineages.

All ASVs in EukBank (Berney et al. 2023) come with information about how they are classified into hierarchical single linkage clusters based on pairwise distances (up to 60 nucleotide differences). Reference sequences from EukRibo are associated to any EukBank ASV cluster containing its closest hit and with a number of differences higher or equal to the number of differences measured with the closest hit. The purpose of this clustering information is to make it possible for any given ASV to identify what the largest ASV cluster is that contains only EukBank ASVs and no EukRibo reference sequence, in order to detect new eukaryotic lineages at various taxonomic levels. We used this clustering information to make sure that we were not missing any ASV of interest, by checking, for all the 1,651 ASVs identified at the previous step, the composition of the largest ASV clusters without reference sequences, which allowed us to add to our selection 487 more ASVs that should be checked for a possible affiliation to our uncharacterized lineages. A script to explore EukBank ASV clusters is available at https://github.com/alijimv/Exploring-Unicellular-Holozoa-Diversity.

We then iteratively repeated these steps. The new selection of 2,138 ASVs was added to our final dataset/alignment of 2,032 sequences as before with the MAFFT v7.525 (Katoh and Standley 2013) option --addfragments. The resulting alignment of 4,170 sequences consisted of 1,719 aligned nucleotide positions after trimming with trimAl v1.5.rev0 (Capella-Gutiérrez et al. 2009) (-gt 0.3). A ML phylogeny was inferred with IQ-TREE v2.3.6 (Minh et al. 2020) as described before and the resulting tree was manually inspected, this time after collapsing branches with UFBoot support <90. We selected all ASVs branching in clades containing some of the environmental 160 sequences but no reference sequences, resulting in 1,470 ASVs. EukBank clustering information was used again to expand this selection a second time to 1,569 ASVs. After re-alignment and trimming done as before (1,759 aligned nucleotide positions), another ML phylogeny was inferred with IQ-TREE v2.3.6 (Minh et al. 2020) as described before. This time 1,184


8

ASVs were retained after visual inspection of the tree; three ASVs branching in uncharacterized lineages but forming extremely long branches were removed. This final pre-selection of ASVs was incorporated again into our final dataset/alignment, which consisted of 3,216 sequences and 1,784 aligned nucleotide positions after trimming. A ML phylogeny was inferred as before, and ASVs affiliated to uncharacterized lineages were selected based on a phylogenetic placement supported by UFBoot values ≥90. This yielded a total of 1,072 ASVs that have been phylogenetically validated as affiliated to our 31 uncharacterized lineages. The maximum-likelihood phylogenetic tree, shown in Fig. 3, and Fig. S2, Newick format and the alignment used to generate it (before and after trimming, FASTA format) are provided in Data S5.

***Relative abundance and geographical and environmental distributions of the uncharacterized lineages***

Relative abundances of characterized versus uncharacterized unicellular Holozoa lineages were estimated using the environmental metadata provided in the EukBank v1.0 (Berney et al. 2023) tables. For characterized lineages we used the “taxogroup1” field in eukbank_18S_V4_asvs.tsv to select ASVs taxonomically annotated as Ichthyosporea, Choanoflagellatea, Filasterea (to which we included *Tunicaraptor*), and Pluriformea (=Corallochytrea). For the uncharacterized lineages identified in this study, we used the affiliated ASVs that we validated phylogenetically before. We then used eukbank_18S_V4_counts.tsv to sum, for all lineages, the read counts (“nreads” field) of their affiliated ASVs in the samples where they are present (“sample” field), separately for each environmental category that the samples belong to (“envplot” field in eukbank_18S_V4_samples.tsv). When an ASV affiliated to one of our uncharacterized lineages was also classified within a described Holozoa lineage based on the EukBank “taxogroup1” annotation, its reads were counted exclusively toward the uncharacterized lineage to avoid double counting (see Data S4).

We also determined the geographical and environmental distributions of the uncharacterized lineages based on the EukBank v1.0 (Berney et al. 2023) environmental metadata. For each lineage, we identified the EukBank samples containing affiliated ASVs and their read counts using the “sample” and “nreads” fields in eukbank_18S_V4_counts.tsv, and obtained their coordinates from the “latitude” and “longitude” fields in eukbank_18S_V4_samples.tsv. The relative abundance of an uncharacterized lineage in each sample was calculated by dividing the total number of reads of the affiliated ASVs by the total number of reads in that sample (“nreads” field in eukbank_18S_V4_samples.tsv). Finally, the environmental distribution of the ASVs affiliated with uncharacterized lineages was estimated by summing read counts across samples within each EukBank environmental category for each ASV. Read counts were then log10(x+1)-transformed (zeros kept as 0) and min-max scaled to values between 0 and 1 within each environmental category, across ASVs (see Data S3 and Data S5).

##### *Data visualization*

Alignments were visualized with AliView v1.28 (Larsson 2014). Trees were visualized with Dendroscope v3.8.10 (Huson and Scornavacca 2012) and FigTree v1.4.4 (Rambaut et al. 2018). The phylogenies in Figures 1, 2, and 3 were visualized and edited in iTOL v7.6 (Letunic and Bork 2024), and those in Figures S1 and S2 in FigTree v1.4.4. Analyses and Figures 4, S3, S4, and S5 were generated in R v4.5.1 (R Core Team 2025) using RStudio v2025.05.1+513 (Posit Team), with the following packages: maps v3.4.3, ggplot2 v3.5.2, dplyr v1.1.4, patchwork v1.3.2, gridExtra v2.3, scales v1.4.0, data.table v1.17.8, and ggforce v0.5.0. All figures were further edited in Inkscape v1.4.2 (Inkscape Project 2025).

Scripts used to explore the relative abundance, and the environmental and biogeographical distribution of EukBank ASVs, and to plot these results, are available at https://github.com/alijimv/Exploring-Unicellular-Holozoa-Diversity.


10

#### Text S2: Descriptions of the Uncharacterized Lineages

**OPI1**

Newly identified lineage within Opisthokonta (Fig. 1 and Fig S1). OPI1 is represented by 20 18S rDNA sequences, retrieved from soil (14), freshwater (5), and freshwater sediment (1) samples (Data S1). It is further supported by 222 affiliated EukBank ASVs (Data S3), found across 842 samples (Data S2) and predominantly associated with terrestrial soil environments: 96.1% in land soil (present in 809 out of 1769 samples), 2.3% in land freshwater (19 out of 35 samples), 0.8% in land sediment (7 out of 132 samples), 0.5% in marine water (4 out of 7928 samples), and 0.4% in marine sediment (3 out 1020 samples). Its highest relative abundance was recorded in a land soil sample (ERR2221528), reaching 3.85%.

**MAOP1**

Previously identified lineage within Holozoa. MAOP1 comprises 19 18S rDNA sequences, obtained from 19 marine water samples (Data S1). It is further corroborated by 315 affiliated EukBank ASVs (Data S3), found across 4,207 samples (Data S2), the vast majority from marine water: 97.4% in marine water (present in 4,097 out of 7928 samples), 1.8% in marine sediment (75 out of 1020 samples), 0.8% in marine organism (32 out of 748 samples), <0.1% in land sediment (1 out of 132 samples), and <0.1% in land freshwater (2 out of 352 samples). Relative abundance was notably high in some samples, reaching a maximum of 7.1548% in a marine water sample (ERR2226849); in 2,635 samples it exceeded 0.01%.

**MAOP2**

Previously identified lineage within Holozoa. MAOP2 is represented by 13 18S rDNA sequences, recovered from marine water (9), marine ice (2), marine sediment (1) and marine organism (1) samples (Data S1). It is further supported by 38 affiliated EukBank ASVs (Data S3), found across 1,578 samples (Data S2) spanning a wide range of habitats, mostly marine: 83.7% in marine water (present in 1,321 out of 7928 samples), 12.0% in marine sediment (189 out of 1020 samples), 1.6% in land sediment (12 out of 132 samples), 1.1% in marine organism (17 out of 748 samples), 1.6% in marine ice (26 out of 95 samples), 0.5% in land soil (8 out of 1769 samples), 0.2% in land freshwater (3 out of 352 samples), and 0.1% in land water (2 out of 42 samples). Relative abundance peaked at 0.9198% in a marine organism sample (ERR2308164).

**HOL1**

Newly identified lineage within Holozoa. HOL1 is represented by 2 18S rDNA sequences, retrieved from soil (1) and freshwater (1) samples (Data S1). It is further supported by 6 affiliated EukBank ASVs (Data S3), found across 34 samples (Data S2) from diverse environments: 58.8% in marine water (present in 20 out of 7928 samples), 32.4% in land soil (11 out of 1769 samples), and 8.8% in land freshwater (3 out of 352 samples). Relative abundance was very low across all samples, with a maximum of 0.0269% recorded in a land soil sample (ERR2233522).

**HOL2**

Newly identified lineage within Holozoa. HOL2 is characterized by 2 18S rDNA sequences, recovered from marine sediment (2) samples (Data S1). It is further represented by 15 affiliated EukBank ASVs (Data S3), found across 132 samples (Data S2) and predominantly associated with marine sediment: 68.2% in marine sediment (present in 90 out of 1020 samples), 27.3% in marine water (36 out of 7928 samples), and 7.6% in marine organism (10 out of 748 samples). Relative abundance peaked at 0.1586% in a marine sediment sample (ERR2308018).

**HOL3**

Newly identified lineage within Holozoa. HOL3 is known from a single 18S rDNA sequence, obtained from a freshwater sediment sample (Data S1). A single affiliated EukBank ASV (Data S3) was found exclusively in land soil (15 out of 1769 samples) (Data S2), with a maximum relative abundance of 0.0116% in sample ERR10038080.

**HOL4**

Newly identified lineage within Holozoa. HOL4 is known from a single 18S rDNA sequence, obtained from a brackish water sample (Data S1). It is complemented by 2 affiliated EukBank ASVs (Data S3), found across 27 samples (Data S2) from diverse environments: 51.9% in land sediment (present in 14 out of 132 samples), 25.9% in marine water (7 out of 7928 samples), 18.5% in land soil (5 out of 1769 samples), and 3.7% in land freshwater (1 out of 352 samples). Relative abundance was low across all samples, with a maximum of 0.0515% recorded in a land soil sample (ERR2232252).

**orphan lineage JN825684**

Newly identified lineage within Holozoa. It is known from a single 18S rDNA sequence, obtained from a freshwater sample (Data S1); no affiliated EukBank ASV was found.



12

**orphan lineage OBEP010278239**

Newly identified lineage within Holozoa. It is represented by two 18S rDNA sequences from groundwater (2) samples (Data S1); no affiliated EukBank ASV was found.

**orphan lineage ORJL011316691**

Newly identified lineage within Holozoa. It is known from a single 18S rDNA sequence from a groundwater sample (Data S1); no affiliated EukBank ASV was found.

**CladeL**

Previously identified lineage within Choanoflagellatea. CladeL is represented by 46 18S rDNA sequences, obtained from soil (10), freshwater (15), freshwater sediment (7), marine water (10), marine sediment (1), marine (2) and saline lagoon (1) samples (Data S1). It is further supported by 66 affiliated EukBank ASVs (Data S3), found across 1,415 samples (Data S2), spanning a broad range of environments: 61.8% in marine water (present in 874 out of 7928 samples), 19.4% in land soil (275 out of 1769 samples), 13.9% in land freshwater (197 out of 352 samples), 1.8% in marine organism (26 out of 748 samples), 1.4% in marine sediment (20 out of 1020 samples), 0.8% in marine ice (12 out of 95 samples), 0.6% in land sediment (8 out of 132 samples), and 0.2% in land organism (3 out of 586 samples). Relative abundance reached a maximum of 1.5861% in a marine water sample (ERR2322065).

**FRESCHO2**

Previously identified lineage within Choanoflagellatea. FRESCHO2 is represented by six 18S rDNA sequences, obtained from freshwater (3), soil (2) and freshwater sediment (1) samples (Data S1). It is further corroborated by 41 affiliated EukBank ASVs (Data S3), found across 282 samples (Data S2), predominantly from land soil: 75.2% in land soil (present in 212 out of 1769 samples), 8.2% in land organism (23 out of 586 samples), 6.7% in land freshwater (19 out of 352 samples), 4.6% in land sediment (13 out of 132 samples), 2.8% in marine organism (8 out of 748 samples), and 2.5% in marine water (7 out of 7928 samples). Relative abundance reached a maximum of 0.4159% in a land soil sample (ERR2233525).

**FRESCHO3**

Previously identified lineage within Choanoflagellatea. FRESCHO3 comprises 46 18S rDNA sequences, obtained from land organisms (26), freshwater (11), marine sediment (5) and freshwater sediment (4) samples (Data S1). It is further supported by 106 affiliated EukBank ASVs (Data S3), found across 480 samples (Data S2), spanning a broad range of environments: 26.7% in land freshwater (present in 128 out of 352 samples), 21.7% in land soil (104 out of 1769 samples), 21.5% in marine sediment (103 out of 1020 samples), 16.7% in marine water (80 out of 7928 samples), 8.8% in land sediment (42 out of 132 samples), 3.5% in marine organism (17 out of 748 samples), 0.6% in marine ice (3 out of 95 samples), 0.4% in land organism (2 out of 586 samples), and 0.2% in land water (1 out of 42 samples). Relative abundance reached a maximum of 8.1219% in a land freshwater sample (ERR2232775).

**FRESCHO4**

Previously identified lineage within Choanoflagellatea. FRESCHO4 is represented by eight 18S rDNA sequences, obtained from freshwater sediment (5), freshwater (2) and soil (1) samples (Data S1). It is further supported by 27 affiliated EukBank ASVs (Data S3), found across 233 samples (Data S2), predominantly from land soil: 64.4% in land soil (present in 150 out of 1769 samples), 27.9% in land freshwater (65 out of 352 samples), 4.3% in marine organism (10 out of 748 samples), 2.1% in marine water (5 out of 7928 samples), 0.9% in land sediment (2 out of 132 samples), and 0.4% in land organism (1 out of 586 samples). Relative abundance reached a maximum of 0.1883% in a land soil sample (ERR2233185).

**MACHO2**

Previously identified lineage within Choanoflagellatea. MACHO2 is represented by five 18S rDNA sequences, obtained from marine water samples (Data S1). It is further supported by 4 affiliated EukBank ASVs (Data S3), found across 38 samples (Data S2): 60.5% in marine water (present in 23 out of 7928 samples) and 39.5% in marine sediment (15 out of 1020 samples). Relative abundance reached a maximum of 0.075% in a marine sediment sample (SRR5753734).

**MACHO4**

Previously identified lineage within Choanoflagellatea. MACHO4 comprises 12 18S rDNA sequences, obtained from marine water (11) and marine (1) samples (Data S1). It is further corroborated by 54 affiliated EukBank ASVs (Data S3), found across 2,521 samples (Data S2), overwhelmingly from marine water: 99.2% in marine water (present in 2,502 out of 7928 samples), 0.6% in marine sediment (14 out of 1020 samples), 0.1% in land freshwater (2 out of 352 samples), 0.1% in land organism (2 out of 586 samples), and <0.1% in marine organism (1 out of 748 samples). Relative abundance reached a maximum of 3.2464% in a marine water sample (SRR7038750).


14

**CHO1**

Newly identified lineage within Choanoflagellatea. CHO1 is characterized by eight 18S rDNA sequences, retrieved from marine water (8) samples (Data S1). It is further corroborated by 54 affiliated EukBank ASVs (Data S3), found across 927 marine samples (Data S2), predominantly marine water: 90.3% in marine water (present in 837 out of 7928 samples), 5.4% in marine organism (50 out of 748 samples), and 4.3% in marine sediment (40 out of 1020 samples). Relative abundance peaked at 1.4075% in a marine water sample (ERR10234174).

**CHO2**

Newly identified lineage within Choanoflagellatea. CHO2 is known from a single 18S rDNA sequence, obtained from a soil sample (Data S1). It is further supported by 6 affiliated EukBank ASVs (Data S3), found across 59 samples (Data S2), mainly from land soil: 67.8% in land soil (present in 40 out of 1769 samples), 15.3% in land freshwater (9 out of 352 samples), 13.6% in marine organism (8 out of 748 samples), and 3.4% in land sediment (2 out of 132 samples). Relative abundance reached a maximum of 0.1671% in a land soil sample (ERR2245037).

**CHO3**

Newly identified lineage within Choanoflagellatea. CHO3 is known from a single 18S rDNA sequence, obtained from a marine sediment sample (Data S1). It is further supported by 6 affiliated EukBank ASVs (Data S3), found across 87 samples (Data S2): 64.4% in marine sediment (present in 56 out of 1020 samples), 25.3% in marine water (22 out of 7928 samples), and 10.3% in land sediment (9 out of 132 samples). Relative abundance reached a maximum of 0.0759% in a marine sediment sample (SRR7034932).

**CHO4**

Newly identified lineage within Choanoflagellatea. CHO4 is represented by 4 18S rDNA sequences, obtained from marine sediment (4) samples (Data S1). It is further corroborated by 5 affiliated EukBank ASVs (Data S3), found across 102 samples (Data S2): 81.4% in marine sediment (present in 83 out of 1020 samples), 17.6% in marine water (18 out of 7928 samples), and 1.0% in marine organism (1 out of 748 samples). Relative abundance reached a maximum of 0.0784% in a marine sediment sample (SRR7035258).



15

**CHO5**

Newly identified lineage within Choanoflagellatea. CHO5 is known from a single 18S rDNA sequence, obtained from a marine water sample (Data S1). It is further supported by 2 affiliated EukBank ASVs (Data S3), found across 113 samples (Data S2): 94.7% in marine water (present in 107 out of 7928 samples) and 5.3% in marine sediment (6 out of 1020 samples). Relative abundance reached a maximum of 0.0196% in a marine water sample (ERR2226623).

**CHO6**

Newly identified lineage within Choanoflagellatea. CHO6 is known from a single 18S rDNA sequence, obtained from a soil sample (Data S1). The single affiliated EukBank ASV was only found in 2 land soil and 1 land sediment samples (Data S3, Data S2). The maximum relative abundance was 0.0081% in a land soil sample (ERR2233515).

**orphan lineage JBGOPN0101245395**

Newly identified lineage within Choanoflagellatea. It is known from a single 18S rDNA sequence from a freshwater sample (Data S1); no affiliated EukBank ASV was found.

**orphan lineage GU290101**

Newly identified lineage within Choanoflagellatea. It is known from a single 18S rDNA sequence from a freshwater sample (Data S1); no affiliated EukBank ASV was found.

**orphan lineage JAKFZD010052931**

Newly identified lineage within Choanoflagellatea. It is known from a single 18S rDNA sequence in a drinking water sample (Data S1); no affiliated EukBank ASV was found.

**FRESHIP1**

Previously identified lineage within Ichthyophonida. FRESHIP1 comprises 25 18S rDNA sequences, obtained from freshwater (18), marine (6), and soil (1) samples (Data S1). It is further corroborated by 41 affiliated EukBank ASVs (Data S3), found across 142 samples (Data S2), mostly from land: 66.2% in land freshwater (present in 94 out of 352 samples), 16.9% in land soil (24 out of 1769 samples), 9.9% in marine water (14 out of 7928 samples), 3.5% in marine sediment (5 out of 1020 samples), 1.4% in marine organism (2 out of 748 samples), 1.4% in land sediment (2 out of 132 samples), and 0.7% in land water (1 out of 42 samples). Relative abundance reached a maximum of 0.3973% in a land freshwater sample (ERR2238562).



16

**FRESHIP2**

Previously identified lineage within Ichthyophonida. FRESHIP2 is represented by 12 18S rDNA sequences, obtained from freshwater (11) and freshwater sediment (1) samples (Data S1). It is further supported by 18 affiliated EukBank ASVs (Data S3), found across 165 samples (Data S2), predominantly from land freshwater: 55.8% in land freshwater (present in 92 out of 352 samples), 24.2% in land soil (40 out of 1769 samples), 18.2% in land sediment (30 out of 132 samples), 1.2% in land water (2 out of 42 samples), and 0.6% in marine water (1 out of 7928 samples). Relative abundance reached a maximum of 9.663% in a land freshwater sample (ERR2225580).

**SOILIP1**

Previously identified lineage within Ichthyophonida. SOILIP1 is represented by two 18S rDNA sequences, obtained from soil (2) samples (Data S1). It is further corroborated by 38 affiliated EukBank ASVs (Data S3), found across 449 samples (Data S2), overwhelmingly from soil: 90.0% in land soil (present in 404 out of 1769 samples), 9.1% in land freshwater (41 out of 352 samples), 0.2% in land organism (1 out of 586 samples), 0.2% in land sediment (1 out of 132 samples), 0.2% in marine sediment (1 out of 1020 samples), and 0.2% in marine water (1 out of 7928 samples). The maximum relative abundance was 0.2343% in a land soil sample (ERR2245098).

**MAIPh1**

Previously identified lineage within Ichthyophonida. MAIPh1 is represented by five 18S rDNA sequences, obtained from freshwater (4) and marine water (1) samples (Data S1). It is further supported by 4 affiliated EukBank ASVs (Data S3), found across 581 samples (Data S2), spanning terrestrial and marine habitats: 44.1% in marine sediment (present in 256 out of 1020 samples), 38.4% in marine water (223 out of 7928 samples), 16.7% in land freshwater (97 out of 352 samples), 0.7% in land sediment (4 out of 132 samples), and 0.2% in land soil (1 out of 1769 samples). The maximum relative abundance was 0.4922% in a land freshwater sample (ERR2225530).

**orphan lineage LC220088**

Previously identified lineage within Ichthyophonida. It is known from a single 18S rDNA sequence from a freshwater sample (Data S1); no affiliated EukBank ASV was found.

**orphan lineage c-2682_conseq_Otu1022**

Previously identified lineage within Ichthyophonida. It is known from a single 18S rDNA sequence from a freshwater sample (Data S1); no affiliated EukBank ASV was found.



17

### Supplementary Figure Legends

See attached files for Figures S1-S5.

**Figure S1. Uncollapsed version of the 18S rDNA phylogeny of Opisthokonta presented in Fig. 1a, revealing 31 uncharacterized lineages.** The tree shown is the maximum-likelihood (ML) topology inferred from an alignment of 2,032 sequences (1,684 aligned nucleotide positions after trimming) under the GTR+FO+I+R10 substitution model and rooted with Breviatea, Apusomonadida, and a closely related environmental lineage. Node labels show ultrafast bootstrap (UFBoot) & transfer bootstrap expectation support values. The nine uncharacterized lineages recovered at Holozoa level plus OPI1 are shown in pink, the six uncharacterized lineages within Ichthyosporea in dark green, and the 15 uncharacterized lineages within Choanoflagellatea in dark blue. See also Data S1-S2.

**Figure S2. Uncollapsed version of the 18S rDNA phylogeny of Opisthokonta presented in Fig. 3, including the 31 uncharacterized lineages and the EukBank ASVs affiliated to them.** The tree shown is the ML topology inferred from an alignment of the same sequences present in Fig. 1a and S1 plus the 1,184 EukBank ASVs that were assigned to the 31 uncharacterized lineages, under the GTR+FO+R10 model of sequence evolution (1,784 aligned nt positions after trimming). The tree is rooted with Breviatea, Apusomonadida, and a closely related environmental lineage. Node labels show UFBoot support values. Colored boxes as in Fig. S1. Uncolored sequences correspond to environmental diversity not meeting our criteria or EukBank ASVs not affiliated with any of the clades of interest. See also Data S3-S5.

**Figure S3. Geographical and environmental distribution of three uncharacterized Holozoa lineages not included in Fig. 4.** Each group is displayed on a separate map (a-c). Circle size is proportional to the relative abundance of reads within each sample, and circle colors indicate the EukBank environmental category. Only samples with relative abundance ≥0.01% are shown. Overlapping circles indicate multiple samples from the same location. See also Data S6.

**Figure S4. Geographical and environmental distribution of four uncharacterized Ichthyosporea lineages not included in Fig. 4.** Each group is displayed on a separate map (a-d). Circle size is proportional to the relative abundance of reads within each sample, and circle colors indicate the EukBank environmental category. Only samples with relative abundance ≥0.01% are shown. Overlapping circles indicate multiple samples from the same location. See also Data S6.

**Figure S5. Geographical and environmental distribution of 11 uncharacterized Ichthyosporea lineages not included in Fig. 4.** Each group is displayed on a separate map (a-k). Circle size is proportional to the relative abundance of reads within each sample, and circle colors indicate the EukBank environmental category. Only samples with relative abundance ≥0.01% are shown. CHO6 is not shown as no sample reached this threshold. Overlapping circles indicate multiple samples from the same location. See also Data S6.


18

#### Supplementary Data Descriptions

See attached files for Data S1-S6.

**Data S1.** Data_S1.xlsx (in relation to Fig. 1a, Fig. 2, and Fig. S1)
List of the 250 environmental 18S rDNA sequences we identified in the databases we screened that fall into the 31 uncharacterized lineages we present in this study, with information about (1) whether the lineage was observed before or is newly identified in this study; (2) the number of sequence clusters, individual sequences, and EukBank ASVs belonging to each lineage; (3) the phylogenetic position of the lineage based on Fig. 1a; and (4) the study and the type of environment the sequences come from.

**Data S2.** Data_S2.zip (in relation to Fig. 1a, Fig. 2, and Fig. S1)
The maximum-likelihood phylogenetic tree shown in Fig. 1a & 2 (in newick format), and the alignment used to generate it (before and after trimming, in fasta format).

**Data S3.** Data_S3.xlsx (in relation to Fig. 1b, Fig. 3, and Fig. S2)
List of all the EukBank ASVs affiliated with each uncharacterized lineage and number of reads they total across all EukBank samples, aggregated by environmental category.

**Data S4.** Data_S4.xlsx (in relation to Fig. 1b)
Total read counts of the EukBank ASVs affiliated with all unicellular Holozoa lineages, aggregated by environmental category. Uncharacterized lineages within Ichthyosporea and Choanoflagellatea are pooled.

**Data S5.** Data_S5.zip (in relation to Fig. 3 and Fig. S2)
The maximum-likelihood phylogenetic tree shown in Fig. 3 (in newick format), and the alignment used to generate it (before and after trimming, in fasta format).

**Data S6.** Data_S6.xlsx (in relation to Fig. 4 and Fig. S3 to S5)
For each uncharacterized lineage, list of all the EukBank samples where affiliated ASVs can be found, with the lineage's total read counts in that sample, the total number of sequenced reads in that sample, and the inferred relative abundance. For each sample the environmental category, latitude and longitude are also provided.


19



20

Katoh K, Standley DM. 2013. MAFFT Multiple Sequence Alignment Software Version 7: Improvements in Performance and Usability. *Mol. Biol. Evol.* 30:772–780.

Kozlov AM, Darriba D, Flouri T, Morel B, Stamatakis A. 2019. RAxML-NG: a fast, scalable and user-friendly tool for maximum likelihood phylogenetic inference.Wren J, editor. *Bioinformatics* 35:4453–4455.

Lanave C, Preparata G, Sacone C, Serio G. 1984. A new method for calculating evolutionary substitution rates. *J. Mol. Evol.* 20:86–93.

Larsson A. 2014. AliView: a fast and lightweight alignment viewer and editor for large datasets. *Bioinformatics* 30:3276–3278.

Latz MAC, Grujcic V, Brugel S, Lycken J, John U, Karlson B, Andersson A, Andersson AF. 2022. Short- and long-read metabarcoding of the eukaryotic rRNA operon: Evaluation of primers and comparison to shotgun metagenomics sequencing. *Mol. Ecol. Resour.* 22:2304–2318.

Lemoine F, Domelevo Entfellner J-B, Wilkinson E, Correia D, Dávila Felipe M, De Oliveira T, Gascuel O. 2018. Renewing Felsenstein's phylogenetic bootstrap in the era of big data. *Nature* 556:452–456.

Letunic I, Bork P. 2024. Interactive Tree of Life (iTOL) v6: recent updates to the phylogenetic tree display and annotation tool. *Nucleic Acids Res.* 52:W78–W82.

Li W, Godzik A. 2006. Cd-hit: a fast program for clustering and comparing large sets of protein or nucleotide sequences. *Bioinformatics* 22:1658–1659.

Minh BQ, Schmidt HA, Chernomor O, Schrempf D, Woodhams MD, Von Haeseler A, Lanfear R. 2020. IQ-TREE 2: New Models and Efficient Methods for Phylogenetic Inference in the Genomic Era.Teeling E, editor. *Mol. Biol. Evol.* 37:1530–1534.

Posit Team. RStudio: Integrated Development Environment for R. Available from: https://posit.co/

R Core Team. 2025. R: A Language and Environment for Statistical Computin. Available from: https://www.R-project.org

Rambaut A, Suchard MA, Xie D, Drummond AJ. 2018. Figtree.

Rognes T, Flouri T, Nichols B, Quince C, Mahé F. 2016. VSEARCH: a versatile open source tool for metagenomics. *PeerJ* 4:e2584.

Sayers EW, Beck J, Bolton EE, Brister JR, Chan J, Connor R, Feldgarden M, Fine AM, Funk K, Hoffman J, et al. 2025. Database resources of the National Center for Biotechnology Information in 2025. *Nucleic Acids Res.* 53:D20–D29.

Schloss PD, Westcott SL, Ryabin T, Hall JR, Hartmann M, Hollister EB, Lesniewski RA, Oakley BB, Parks DH, Robinson CJ, et al. 2009. Introducing mothur: Open-Source, Platform-Independent, Community-Supported Software for Describing and Comparing Microbial Communities. *Appl. Environ. Microbiol.* 75:7537–7541.

Seemann T. 2013. tseemann/barrnap. Available from: https://github.com/tseemann/barrnap

Shen W, Sipos B, Zhao L. 2024. SeqKit2: A Swiss army knife for sequence and alignment processing. *iMeta* 3:e191.

Tavaré S. 1986. Some probabilistic and statistical problems on the analysis of DNA sequences. In: Some Mathematical Questions in Biology: DNA Sequence Analysis. Vol. 17. Lec. Math. Life Sci. Providence, RI: American Mathematical Society. p. 57–86.

Urrutia A, Mitsi K, Foster R, Ross S, Carr M, Ward GM, Van Aerle R, Marigomez I, Leger MM, Ruiz-Trillo I, et al. 2022. *Txikispora philomaios* n. sp., n. g., a micro -eukaryotic pathogen of amphipods, reveals parasitism and hidden diversity in Class Filasterea. *J. Eukaryot. Microbiol.* 69:e12875.



22
