## Supplementary figures and images for "Uncharacterized unicellular relatives reveal key gaps in sampling diversity for reconstructing animal origins"

### Figure S1

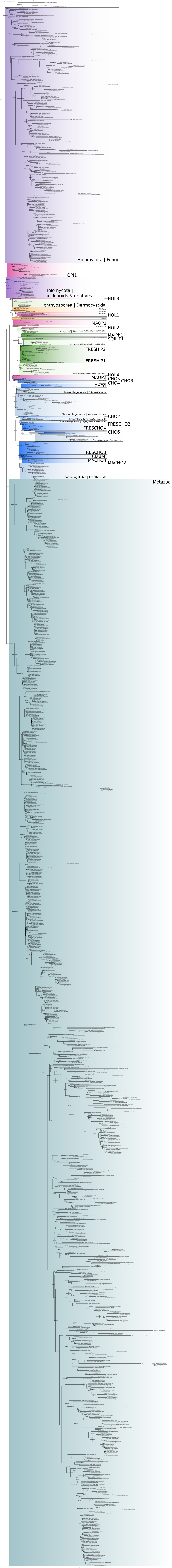

### Figure S2

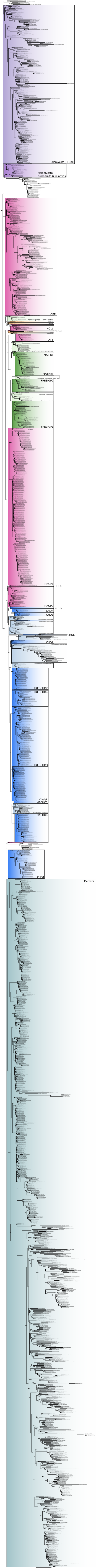

### Figure S3

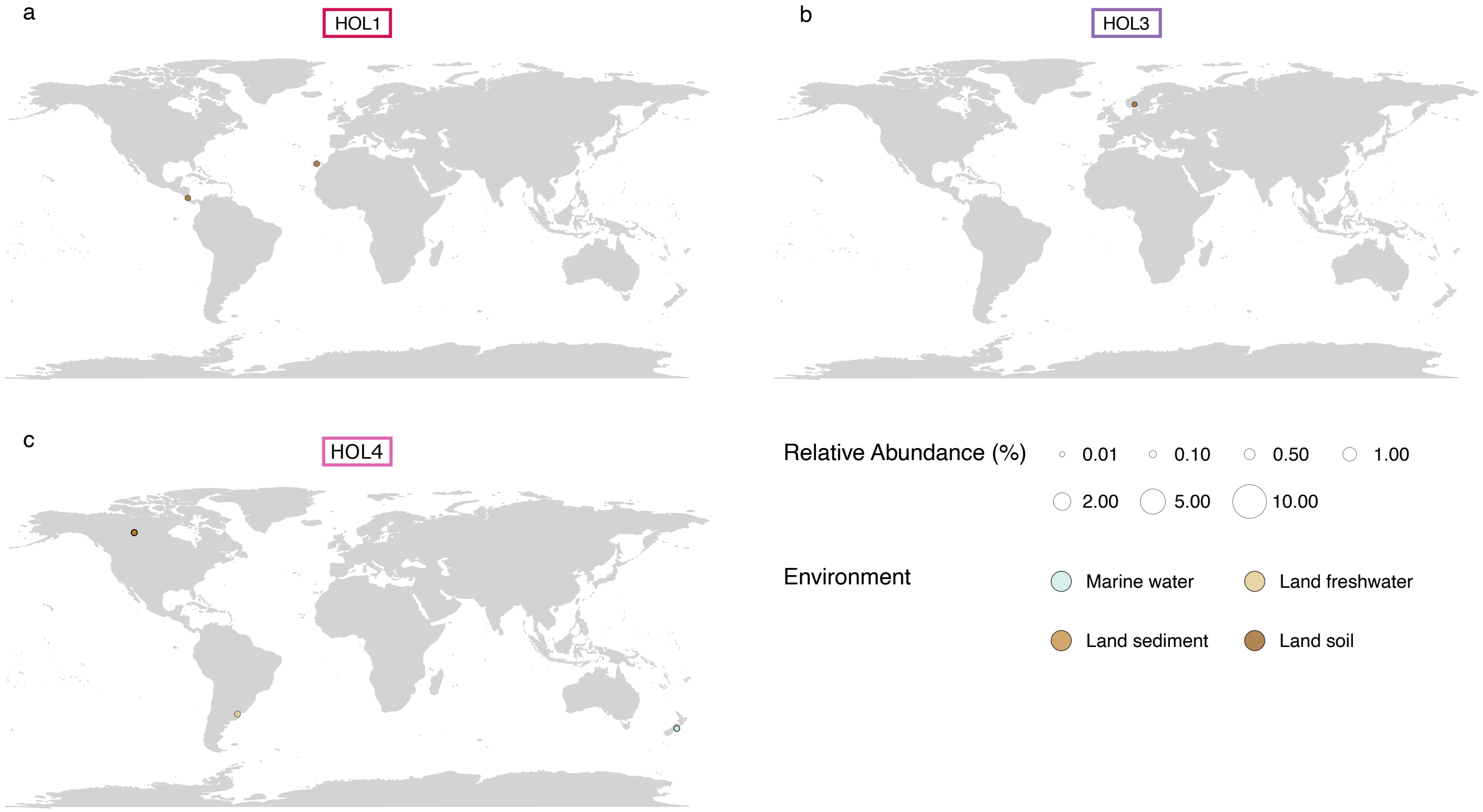

### Figure S4

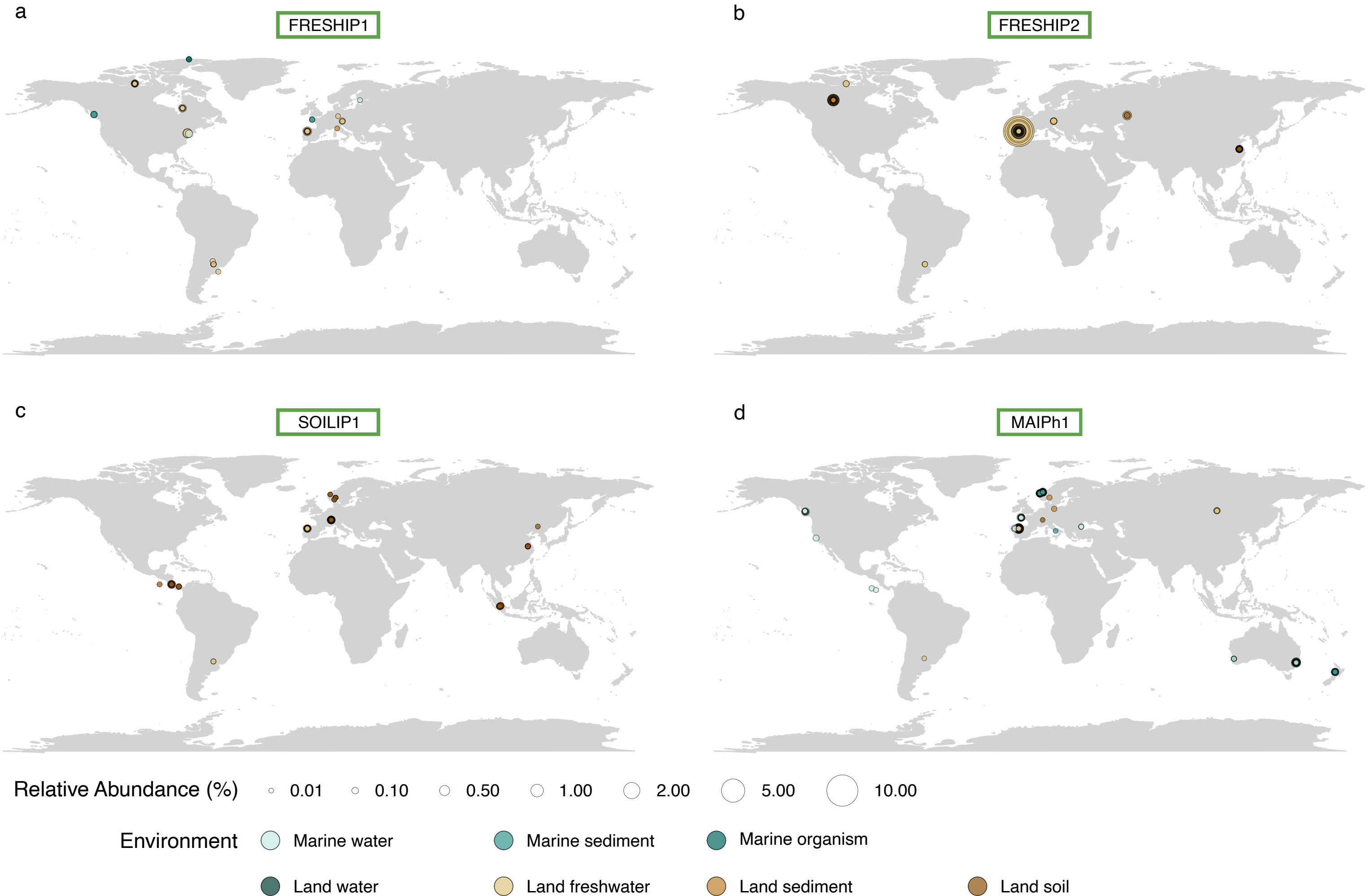

### Figure S5

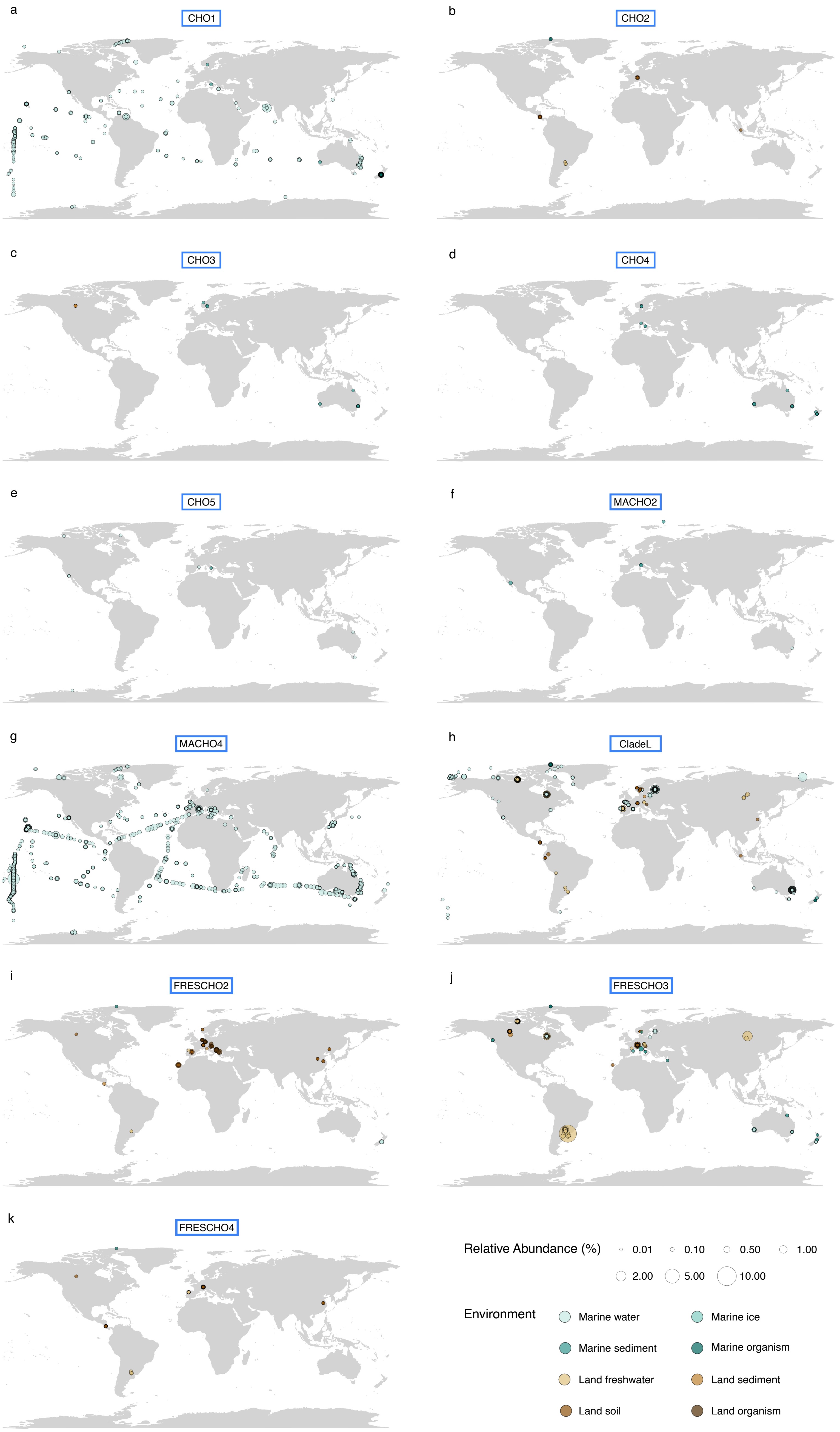
